# Glycolysis/gluconeogenesis specialization in microbes is driven by biochemical constraints of flux sensing

**DOI:** 10.1101/2021.05.07.443112

**Authors:** Severin Josef Schink, Dimitris Christodoulou, Avik Mukherjee, Edward Athaide, Viktoria Brunner, Tobias Fuhrer, Gary Andrew Bradshaw, Uwe Sauer, Markus Basan

## Abstract

Central carbon metabolism is highly conserved across microbial species, but can catalyze very different pathways depending on the organism and their ecological niche. Here, we study the dynamic re-organization of central metabolism after switches between the two major opposing pathway configurations of central carbon metabolism, glycolysis and gluconeogenesis in *Escherichia coli, Pseudomonas aeruginosa* and *Pseudomonas putida*. We combined growth dynamics and dynamic changes of intracellular metabolite levels with a coarse-grained model that integrates fluxes, regulation, protein synthesis and growth and uncovered fundamental limitations of the regulatory network: after nutrient shifts, metabolite concentrations collapse to their equilibrium, rendering the cell unable to sense which direction the flux is supposed to flow through the metabolic network. The cell can partially alleviate this by picking a preferred direction of regulation at the expense of increasing lag times in the opposite direction. Moreover, decreasing both lag times simultaneously comes at the cost of reduced growth rate or higher futile cycling between metabolic enzymes. These three trade-offs can explain why microorganisms specialize for either glycolytic or gluconeogenic substrates and can help elucidate the complex growth patterns exhibited by different microbial species.

**Graphical synopsis:** 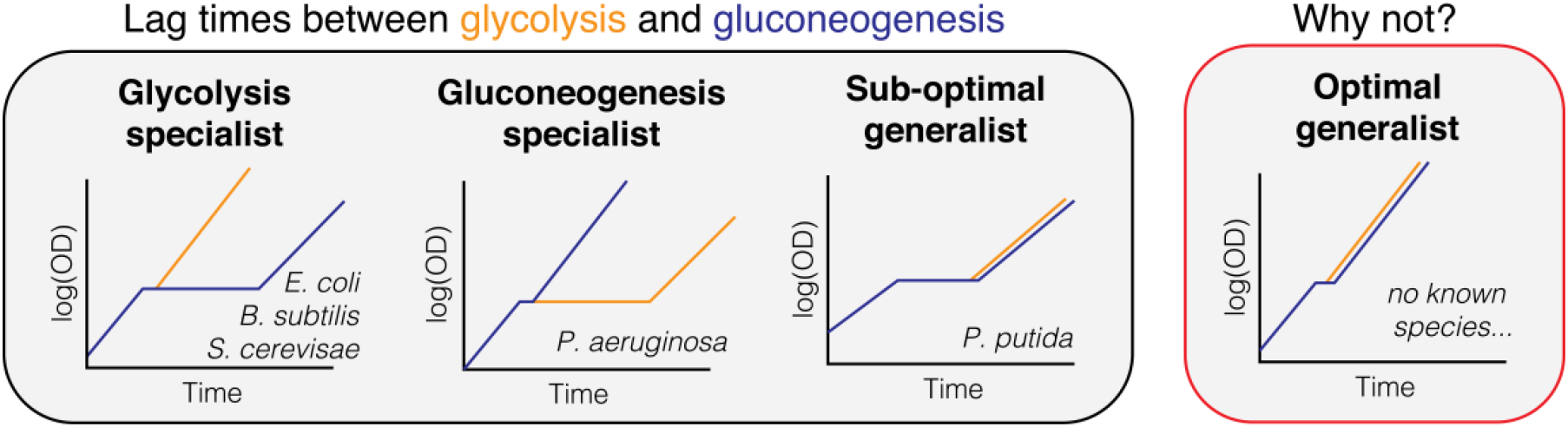

**Standfirst text:** Microbes face a series of fundamental trade-offs that limit their ability to optimize simultaneously for both glycolytic and gluconeogenic growth.

**Bullet points:**

- Lag times between glycolysis and gluconeogenesis show asymmetry in many microbes: A long lag in one direction, but a short lag in the other.
- Long lag times are caused by an inability to sense fluxes after nutrient shifts.
- With existing regulation, lag time asymmetry can only be overcome by reducing either growth rate or increasing futile cycling in metabolism.

## Introduction

Whether in nature, microbiomes or infections, microbes frequently encounter changing environments (Hardcastle and Mann, 1968; Fenchel, 2002; Stocker, 2012; Battin et al., 2016; Forsyth et al., 2018) and their ability to adapt quickly is a key determinants of fitness. But in comparison with steady state exponential growth, understanding of the physiology of growth transitions, in particular what sets the time-scales of adaptation, has remained largely elusive. For steady state exponential growth, metabolic models have made substantial progress over the last two decades, elucidating the flux and regulatory networks that govern the coordination of microbial metabolism (Bennett et al., 2009; Bordbar et al., 2014; Chubukov et al., 2014; Gerosa et al., 2015a; Link et al., 2013; Noor et al., 2010, 2014; Vasilakou et al., 2016). Such metabolic models were successfully expanded to dynamic environments (Zampar et al., 2013; Chassagnole et al., 2002; Chakrabarti et al., 2013; Saa and Nielsen, 2015; Andreozzi et al., 2016; Yang et al., 2019) and used to gather kinetic information about metabolism, using perturbations (Link et al., 2013), stimulus response experiments (Chassagnole et al., 2002) or sequential nutrient depletion (Yang et al., 2019) to validate and improve metabolic models. But, dynamic changes of metabolism like shifts in growth conditions continue to pose a considerable challenge, and it is still unclear what determines how long bacteria need to adapt upon a change of the environment.

One example of such a switch happens when microbes deplete their primary nutrient. *Escherichia coli* preferentially utilizes hexose sugars like glucose that are metabolized via glycolysis (Gerosa et al., 2015b). To maximize growth on sugars, *E. coli* excretes substantial ‘overflow’ production of acetate, even in the presence of oxygen (Basan et al., 2015a, 2017). This naturally leads to bi-phasic growth, if no other microbe is around to utilize this bi-product, where initial utilization of glucose is followed by a switch to acetate. Similar growth transitions from preferred glycolytic substrates to alcohols and organic acids ubiquitously occur for microbes in natural environments (Buescher et al., 2012; Otterstedt et al., 2004; Zampar et al., 2013). Since these fermentation products are all gluconeogenic, they require a reversal of the flux direction in the glycolysis pathway, which results in multi-hour lag phases caused by the depletion of metabolite pools throughout the gluconeogenesis pathway (Basan et al., 2020). Similar long lag times in glycolytic to gluconeogenic shifts were observed for *Bacillus subtilis* and the yeast *Saccharomyces cerevisiae* (Basan et al., 2020). Shifts in the opposite direction, however, from gluconeogenic substrates to glycolytic ones, occur much more quickly in *E. coli* and other preferentially hexose fermenting microbes, in some cases even without detectable lag phases (Basan et al., 2020).

In our previous work (Basan et al., 2020), we showed how the growth rate dependence of enzyme expression leads to a universal relation between lag times and preshift growth rates and found evidence that futile cycling at irreversible metabolic reactions plays an important role for causing lag times. However, we were unable to answer the most fundamental questions raised by these observations: Why are microorganisms like *E. coli* or *S. cerevisiae* unable to overcome lag phases by expressing more metabolic enzymes or allosteric regulations that turn off futile cycling after metabolic shifts? Given the small number of enzymes involved in these irreversible reactions, their cost in terms of proteome allocation is likely minimal. Instead, microbes like *E. coli* appear to be intentionally limiting enzyme expression and decreasing their growth rates on many glycolytic substrates (Basan et al., 2017). Moreover, why do shifts from glycolytic to gluconeogenic conditions result in lag times of many hours, while shifts from gluconeogenic to glycolytic conditions only take minutes? Given the symmetry of central metabolism, one would expect similar lag phases in the opposite direction. Is this preference for glycolysis a fundamental property of central metabolism or rather an evolutionary choice of individual species? At the core of these questions is a gap in understanding of how central carbon metabolism adjusts itself to nutritional changes.

Here, we study growth and metabolite dynamics of *E. coli, Pseudomonas aeruginosa* and *Pseudomonas putida* using a kinetic model of central carbon metabolism to overcome this challenge. Our model coarse-grains central metabolism to a low number of irreversible and reversible reactions, which allows us to focus on the dynamics of key metabolites and their regulatory action. The model couples metabolism to enzyme abundance via allosteric regulation and enzyme expression to the concentration of regulatory metabolites via transcriptional regulation and flux dependent protein synthesis. Our formulation of metabolism and growth bridges fast metabolic time scales with slow protein synthesis. As we demonstrate, our model can explain a major reorganization of metabolism in response to nutrients shifts: the switching of the directionality of metabolic flux between glycolysis and gluconeogenesis. Dependent on the required directionality of flux in central metabolism, enzymes catalyzing the required flux direction are expressed and catalytically active, while enzymes catalyzing the opposite flux are expressed at low levels and their activities are repressed by allosteric regulation. This self-organization is key for enabling fast growth and preventing costly futile cycling between metabolic reactions in opposing directions, which can inhibit flux and deplete ATP in the process.

Reestablishing this self-organization after growth shifts is limited by biochemical constraints to sense fluxes and to regulate accordingly. When metabolite levels transiently collapse, allosteric and transcriptional regulation cannot distinguish between glycolysis and gluconeogenesis, rendering the cell unable to sense to the direction of flux. By choosing the activity of metabolic enzymes at these low metabolite levels to favor one direction, the cell can enable fast switching at the expense of the other direction. This choice of direction at low metabolite concentrations becomes the ‘default state’ of central metabolism and determines the substrate preference.

According to the model, the preferred direction does not need to be glycolysis, and in principle gluconeogenic specialists with a gluconeogenic ‘default’ state could have evolved, too. Indeed, we showed that *P. aeruginosa* shows reversed lag time and growth phenotypes compared to those of *E. coli*, which verified that long lag times to glycolytic substrates are caused by the same inability to sense flux after nutrient shifts.

## Results

### An integrated, self-consistent kinetic model of glycolysis / gluconeogenesis

In a shift between glycolysis and gluconeogenesis, flux in central metabolism needs to be reversed. To understand what limits the speed of adaptation between those two modes of flux, we turn to a theoretical model of central metabolism. But because the complexity of central metabolism with intertwined regulation at different levels prevents tracing quantitative phenotypes to their molecular origins, we sought to focus on the biochemical pathway topology with its key regulations that differentiate glycolysis and gluconeogenesis and constructed a minimal model of central metabolism. The model is illustrated in Box 1 and described in detail in the SI. It is based on topology of the biochemical network, the allosteric and the transcriptional regulation of the key the metabolic proteins of *E. coli*, all of which have been well characterized (Berger and Evans, 1991; Ramseier et al., 1995; Johnson and Reinhart, 1997; Pham and Reinhart, 2001; Kelley-Loughnane et al., 2002; Hines et al., 2006; Fenton and Reinhart, 2009).

#### Box 1 Integrated kinetic model of central carbon metabolism.

(**A**) Detailed metabolic reaction network and (B) minimal network of central carbon metabolism. Coarse-graining was done by combining irreversible glycolytic (orange) and gluconeogenic reactions (blue), as well as metabolites. Influx can either occur from glycolytic carbon sources (e.g. glucose) or gluconeogenic carbon sources, (e.g. tricarboxylic acid (TCA) cycle carbon like acetate or malate). (1) Gatekeepers to the central section of glycolysis and gluconeogenesis are the two irreversible reactions (gly^up^, gng^up^ and gly^low^, gng^low^) that feed and drain FBP and PEP. The irreversible reactions are allosterically regulated by FBP (Fructose 1-6-bisphosphate) and PEP (phosphoenolpyruvate), where ‘outward’ facing reactions are activated (green arrows) and ‘inward’ facing reactions are repressed (red arrow). (2) Biomass production requires precursors from glycolytic carbons, PEP and gluconeogenic carbons (i.e., from TCA cycle). (3) Glycolytic and gluconeogenic enzymes are regulated by Cra, which is in turn modulated by FBP. (**C**) Mathematical formulation of the model. Numbers correspond to features in panel B. (1) Fluxes r_i_ of enzymes i depend on enzyme abundances ϕ_i_, catalytic rates k_cat,i_ and allosteric regulations, modeled as a Hill function below its maximal saturation 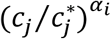, where c_j_ is the concentration of the regulatory metabolite and 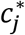 is a reference concentration. Reversible fluxes are modeled with simple mass action kinetics. (2) Biomass production is implemented in the model as single reaction that drains all three metabolites simultaneously at catalytic rate k_cat,BM_. (3) Enzyme expression depend linearly on FBP concentration c_FBP_. Growth rate: μ, steady state abundance: 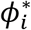, steady state concentration 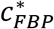 and x_i_ & x_j_ modulate the sensitivity of regulation to FBP. Glycolytic and gluconeogenic enzymes are produced as part of protein synthesis. Thus in the model, flux through metabolism automatically leads to synthesis of metabolic enzymes and biomass production, resulting in dilution of existing enzymes.

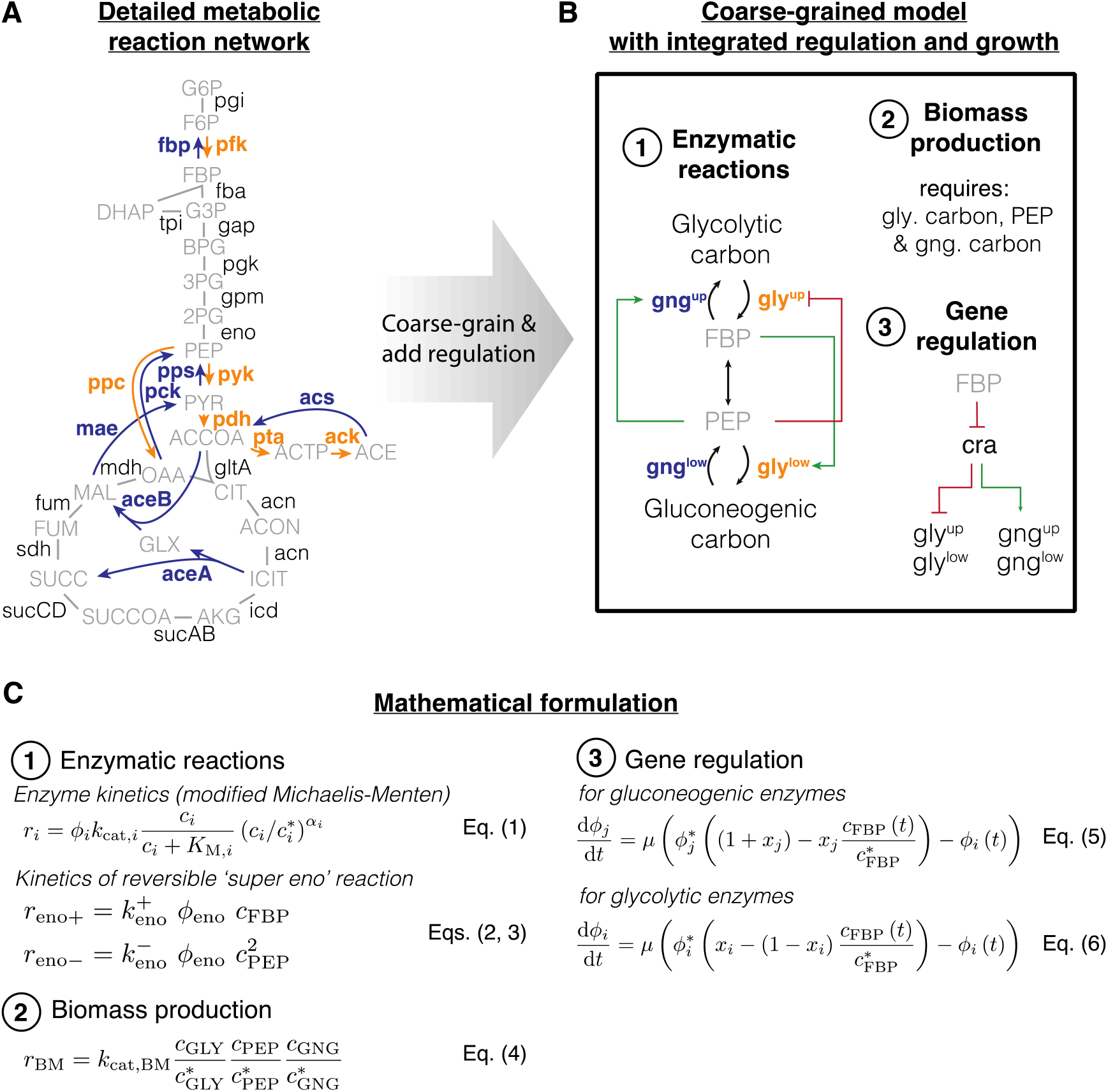

The defining feature of the model is a coarse-graining of the irreversible reactions (one-directional arrows in ‘orange’ and ‘blue’, Box 1A) in the upper and lower part of central metabolism into single irreversible reactions (one-directional ‘black’ arrows in Box 1B). While not irreversible in an absolute sense, so-called irreversible reactions are thermodynamically favored so much in one direction that they can be effectively considered as irreversible (Noor et al., 2014). As a result, these irreversible reactions in central metabolism need to be catalyzed by distinct enzymes that perform distinct reactions For example, Fructose 6-phosphate (F6P) is converted to Fructose 1-6-bisphosphate (FBP) by enzyme PfkA using ATP. The opposite direction, FBP to F6P, is performed by a different enzyme, Fbp, which splits off a phosphate by hydrolysis. Each of the two reactions follows a free energy gradient and are irreversible. If both enzymes are present and active then the metabolites will be continuously interconverted between F6P and FBP and in each interconversion one ATP is hydrolyzed to ADP and phosphate. This is a ‘futile cycle’. It drains the cell’s ATP resource and prevents flux going through the biochemical network. Because of this importance of irreversible reactions and futile cycling, we implement irreversible enzymes (‘bold font, blue/orange’ in Box 1A&B) and their allosteric regulation (‘green’ and ‘red’ arrows in Box 1B) in the model. To successfully switch flux directions, the cell needs to express irreversible enzymes in the new direction, up-regulate their activity and repress enzyme activity in the opposing direction. Uptake of carbons from the environment is modeled as a flux to the substrates of the irreversible reactions, either glycolytic or gluconeogenic carbons depending on the availability.

The metabolites ‘sandwiched’ between the irreversible reactions are coarse-grained into the first and last metabolites of the series of reversible reactions connecting the irreversible reactions, FBP and PEP (phosphoenolpyruvate). These metabolites regulate the activity and expression of the irreversible enzymes (Box 1B and SI Sec. 2).

In total, the model encompasses four irreversible reactions, each regulated allosterically by either FBP or PEP, and transcriptionally by FBP via Cra, and one reversible reaction that connects FBP and PEP. We used measured metabolite concentrations for growth on glucose (Kochanowski et al., 2013a) and Michaelis constants (Berman and Cohn, 1970; Zheng and Kemp, 1995; Donahvue et al., 2000) to constrain enzymatic parameters and biomass yield (Link et al., 2008) and density (Basan et al., 2015b) on glucose to constrain fluxes (SI Sec. 4). We used the level of futile cycling in the upper and lower reactions in exponential glucose growth, which summarize the effect of enzyme abundance and allosteric regulation, as fitting parameters such that the model reproduces the observed lag times in this paper; see SI Sec. 4.2 for details.

While the model in Box 1 was formulated to coarse-grain glycolysis via the Embden-Meyerhof-Parnas (EMP) pathway, the dominant glycolytic pathway of *E. coli* growing on glucose (Gerosa et al., 2015a), other glycolytic pathways, such as the Entner-Doudoroff (ED) or pentose phosphate pathway (PPP), have a similar topology. In ED glycolysis, phosphogluconate dehydratase (Edd) and KDPG aldolase (Eda) are irreversible reactions that feed into the chain of reversible reactions, analogous to 6-phosphofructokinase (pfk) in the EMP pathway. The coarse-grained model thus should capture these alternative pathways too.

### Central carbon metabolism self-organizes in response to substrate availability

To test whether this simple model could recapitulate steady state glycolytic and gluconeogenic growth conditions, we calibrated it with published metabolite and proteomics data of *E. coli*, which is well-characterized in steady state exponential growth on glucose and acetate as sole carbon substrates (Basan et al., 2020). Indeed, the model reached distinct steady states for glycolytic and gluconeogenic conditions, which we summarized graphically with font size indicating enzyme and metabolite abundance and line widths indicating the magnitude of fluxes (Fig. 1A&B). Active regulation is shown in colored lines, while inactive regulation are grey, dashed lines. We quantitatively compare enzyme and metabolite abundances to experimental measurements in Fig. 1C-E and find that the coarse-grained model can describe the reorganization of metabolism well, despite the simplifications of the metabolic and regulatory networks.

**Figure 1.**
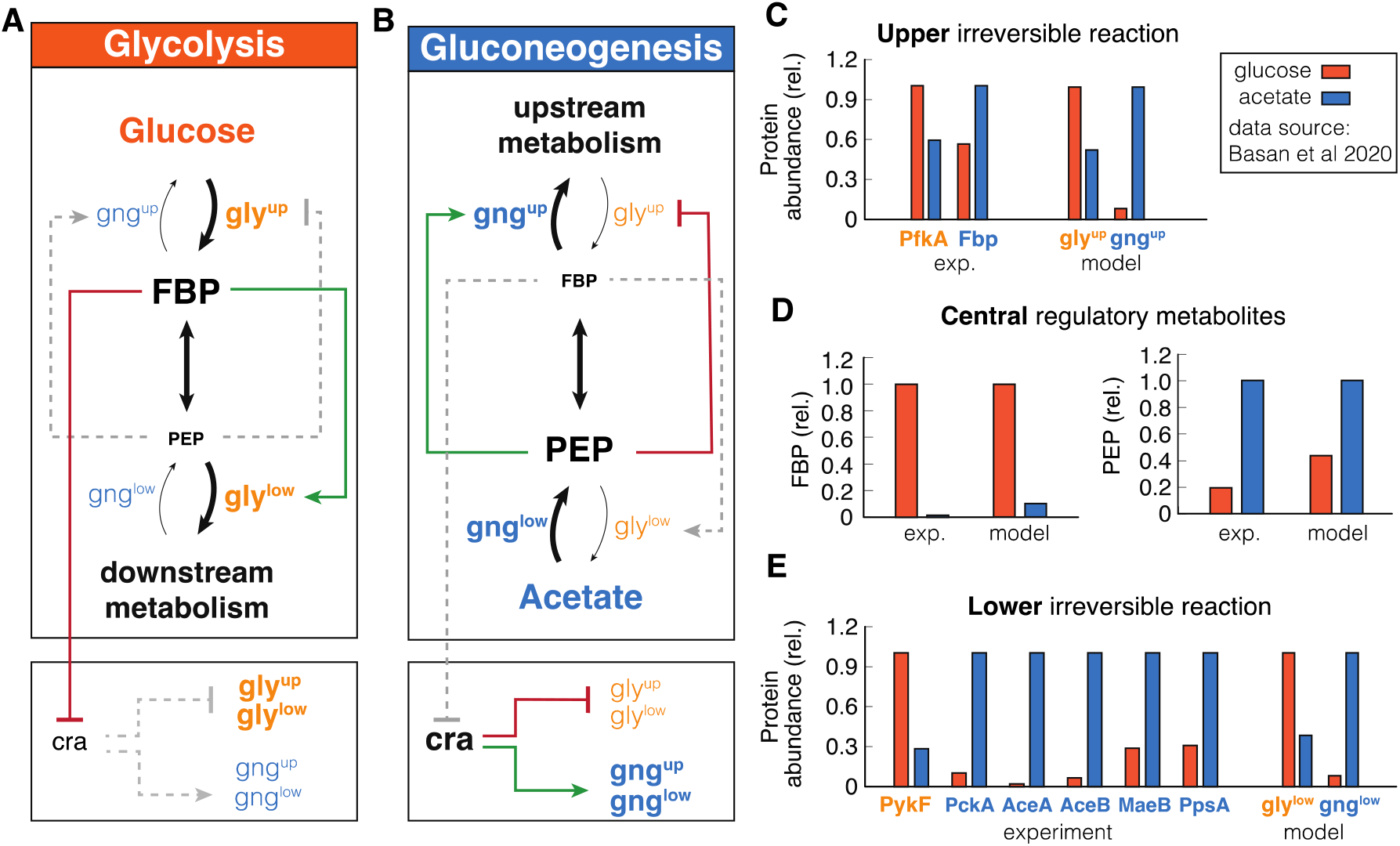
Self-organization of metabolism in glycolysis and gluconeogenesis. (**A&B**) Graphic summary of the reorganization in glycolysis and gluconeogenesis. Linewidth of reactions arrows indicate magnitude of flux. Font size of metabolites and enzymes indicate metabolite concentrations and enzyme abundances, respectively. Active regulation is indicated by red/green color, inactive regulation is grey and dashed. (**C, D**&**E**) Calibration of model to experimental data (from (Basan et al., 2020)) metabolite concentrations and enzyme abundances relative to the highest concentration or abundance. Note the striking, differential regulation of FBP and PEP, high in one condition and low in the other.

The simulation helps to understand how central metabolism self-organizes in glycolytic and gluconeogenic conditions and how allosteric and transcriptional regulation optimize fluxes and minimize futile cycling during exponential growth. As shown in Fig. 1C, in ‘orange’, during glycolytic conditions, the simulation reached a steady state with high FBP levels and low PEP levels. As illustrated in Fig. 1A, the high FBP pool activates lower glycolysis, while the low PEP pool derepresses upper glycolysis and deactivates upper gluconeogenesis. This suppression of gluconeogenic fluxes in glycolysis reduces futile cycling, i.e., circular fluxes at the irreversible reactions, thereby streamlining metabolism. On a transcriptional level, the high FBP pool represses Cra, which in turn derepresses the expression of glycolytic enzymes and inhibits the expression of gluconeogenic enzymes. This results in high levels of glycolytic enzymes and low levels of gluconeogenic enzymes in the simulation (Fig. 1D&E, right panels).

In gluconeogenic conditions (‘blue’ in Fig. 1), we find precisely the complementary configuration of central carbon metabolism. Simulation and experiments show low FBP and high PEP pools (Fig. 1C). As illustrated in Fig. 1B, high PEP represses upper glycolysis and activates upper gluconeogenesis, while low FBP deactivates lower glycolysis. Low FBP also derepresses Cra, which leads to high expression of gluconeogenic enzymes and low expression of glycolytic enzymes (Fig. 1D, right panels).

Next we tested if the model could recapitulate how varying growth rates on glycolytic and gluconeogenic nutrients affects metabolite levels and protein expression in *E. coli* (Gerosa et al., 2015b; Hui et al., 2015). In particular, it has been shown experimentally that FBP acts like a flux sensor and FBP concentration linearly increases with glycolytic flux (EV Fig. 1A) (Kochanowski et al., 2013b), which is recapitulated by our simulation (EV Fig. 1D), under the condition that enzymes catalyzing the reversible reaction are far from saturation. The linear increase of FBP concentration with growth rate results in a linear growth rate dependence of gluconeogenic and glycolytic enzyme abundances in the simulation, in good agreement with experimental measurements of enzyme abundances from proteomics (EV Fig. 1 compare B&C with E&F) (Hui et al., 2015). Together, these results show that integrating the transcriptional and allosteric regulation of FBP and PEP in the coarse-grained model suffices to describe the major re-configuration of central metabolism in glycolysis and gluconeogenesis.

### Central carbon metabolism of E. coli is primed for switches to glycolysis

Equipped with this model, we next address the mechanistic basis for the extended lag phases of *E. coli* upon nutrient shifts from glycolytic to gluconeogenic conditions. When shifted from glucose to acetate *E. coli* shows a lag time with almost no growth for around 5h (Fig. 2A, data: (Basan et al., 2020)). We can reproduce this lag with our model (Fig. 2B, Appendix Fig. S1-5) when we fit pre-shift futile cycling, which is a measure for enzyme abundances and allosteric regulations; see SI Sec. 2 for details. All model solutions for *E. coli* shown in this paper are generated with the parameters generated from this fit. The model captures the slow adaptation of glycolytic and gluconeogenic enzymes, the major change of which occurs only towards the end of the lag phase (Appendix Fig. S6). Investigating the origin of the growth arrest in the simulation, we found that during lag phase, the concentrations of upper glycolytic precursors (which includes Fructose 6-phosphate (F6P) and Glucose 6-phosphate (G6P)) remained very low compared to their steady state values, which matches published experimental evidence of F6P measurements (Basan et al., 2020) (Fig. simulation: 2C, data 2D). This indicates that essential precursors are limited, and thereby, according to Eq. (4) growth rate during lag phase stalls.

**Figure 2.**
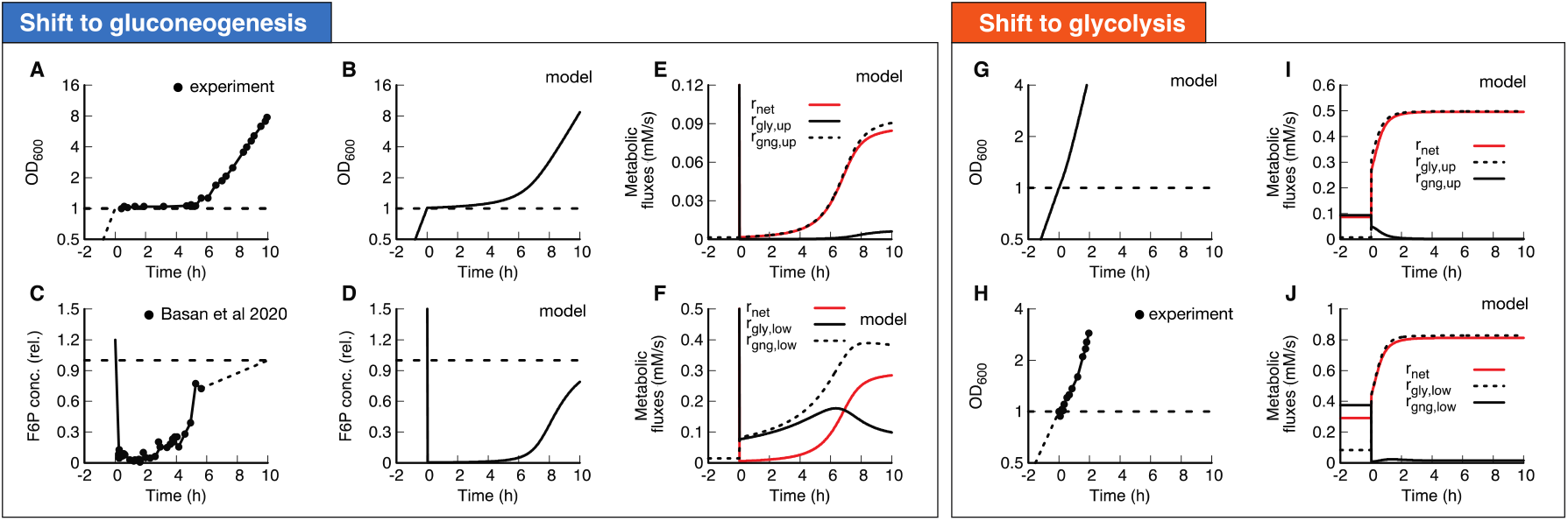
Shifts between glycolysis and gluconeogenesis. (**A**) Experimental and (**B**) model of optical density after shift of E. coli from glucose to acetate. Growth shows a substantial lag before it recovers. (**C**) Experimental and (**D**) model of F6P (normalized to the final state) collapses after shit to acetate, and continues to stay low throughout lag phase. Because F6P is an essential precursor for biomass production, this limitation effectively stops biomass growth. Data points shows a single time-series from (Basan et al., 2020). (**E**&**F**) Fluxes of all irreversible reactions in units of intracellular concentration per time. Especially fluxes in lower glycolysis/gluconeogenesis are of equal magnitude, leading to a futile cycle, where no net flux (red line) through central carbon metabolism can be established. (**G-J**) Optical density and metabolic fluxes for the reversed shift from acetate to glucose shows immediate growth and no intermittent futile cycling. The dynamics of all enzyme abundances, regulation and fluxes for both shifts are shown in Appendix Fig. S1-5 in detail. The model also correctly predicts that enzyme abundances only adapt late in the lag phase (Appendix Fig. S6).

In the simulation, the F6P limitation is caused by low net fluxes in upper and lower gluconeogenesis (Fig. 2E&F, red lines). Previously, it was suggested that futile cycling between gluconeogenic and glycolytic enzymes could contribute to this flux limitation (Basan et al., 2020), supported by the observation that overexpression of glycolytic enzymes in upper or lower glycolysis strongly impaired switching and resulted in much longer lag times (Basan et al., 2020). The simulation allows us to probe the effect of futile cycling *in silico*, which cannot be directly measured experimentally. Indeed, we found for our default *E. coli* parameters that residual lower glycolytic flux almost completely canceled the flux from gluconeogenesis, i.e., 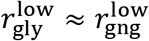 (solid and dashed black lines in Fig. 3F), such that net gluconeogenic flux remained close to zero (red line, Fig. 2E&F). Thus, this futile cycling appears to be the main reason for limiting net flux throughout the lag phase.

**Figure 3.**
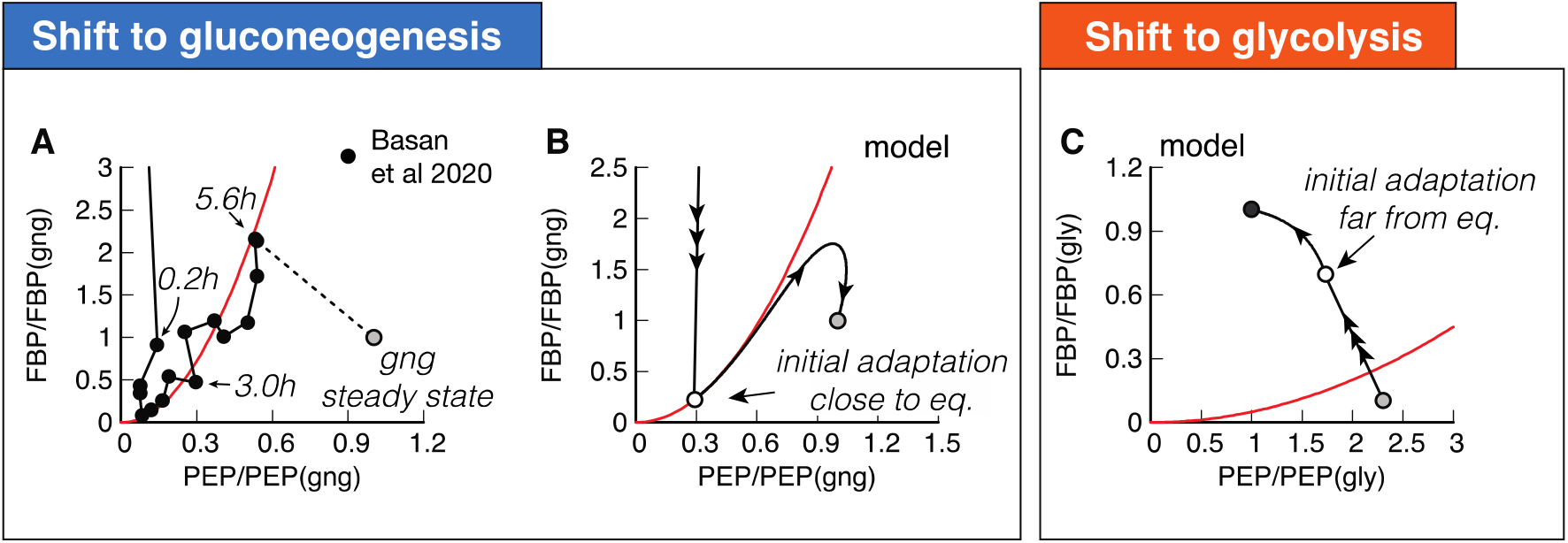
Molecular cause for asymmetric recovery dynamics in E. coli. (**A**) Recovery of FBP and PEP of after a shift from glucose to acetate, shows a distinctive joint increase, followed by an overshoot of FBP. Red line is a quadratic guide to the eye. Final acetate steady state is drawn as grey symbol and used to normalize both FBP and PEP levels. Data is a single time-series from (Basan et al., 2020). (**B**) Model solution of FBP and PEP. After the fast collapse of metabolite levels (triple arrow to white circle), the dynamics closely follows the quadratic FBP-PEP equilibrium Eq. (7). Eventually recovery will diverge away from the equilibrium line, towards the non-equilibrium steady states of gluconeogenesis (grey circle) (**C**) For a shift to glycolysis, metabolite levels do not collapse, but instead land already far from equilibrium (triple arrow to white circle), such that flux is immediately established, and recovery is quick.

The biochemical network and regulation are almost completely symmetric with respect to the direction of flux, so one might naively expect a shift from gluconeogenesis to glycolysis to also result in a long lag. However, experimentally the shift in the opposite direction from gluconeogenesis to glycolysis occurs very quickly in *E. coli* (Fig. 2G) (Basan et al., 2020). Our simulations with the standard *E. coli* parameters can recapitulate that central metabolism adjusted very quickly and growth resumed without a substantial lag phase (Fig. 2H). In striking contrast to the shift to gluconeogenesis, futile cycling played no role in the shift to glycolysis, because both upper and lower glycolytic fluxes got repressed immediately after the shift (Fig. 2I-J, solid black line), such that net flux could build up (Fig. 2I-J, red line). The absence of transient futile cycling, despite the symmetry of regulation and metabolic reactions, means that, according to the model, it must be the allosteric and transcriptional regulations that ‘prime’ central metabolism of *E. coli* for the glycolytic direction.

### Molecular cause of preferential directionality

To understand the molecular cause of the asymmetric response and lag phases, we investigated the role of allosteric and transcriptional regulation in our simulation. During steady state growth, the differential regulation during glycolysis and gluconeogenesis is achieved by PEP and FBP, the metabolites that are “sandwiched” between the two irreversible reactions and connected by a series of reversible enzymes, coarse-grained in our model into the ‘super-eno’ enzyme. First, we focused on regulation during exponential growth and wanted to investigate how the cell achieves differential regulation of glycolytic and gluconeogenic enzymes using the metabolites FBP and PEP. In equilibrium, forward and backward reactions would balance, i.e., *r*_ENO+_ = *r*_ENO−_, and no net flux could run through central metabolism, meaning that the cell could not grow. Using Eqs. (2&3), the balance of forward and backward fluxes results in a fixed quadratic dependence of FBP and PEP in equilibrium,

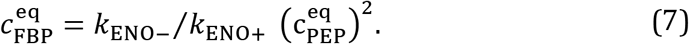

The form of Eq. (7) is specific to F6P converting to FBP being the irreversible step of upper glycolysis and can change if pathways such as Entner-Doudoroff (ED) or pentose phosphate pathway (PPP) are dominant.

Close to the equilibrium, FBP and PEP levels go up and down together, rather than the opposing directions, as observed for glycolytic and gluconeogenic growth (Fig. 1A&B).

This results in low net flux and very slow growth. Hence, for steady state growth, the equilibrium must be broken and FBP ≫ PEP or FBP ≪ PEP, such that either glycolytic flux is bigger than gluconeogenic or vice-versa (*r*_ENO+_ ≫ *r*_ENO-_ and *r*_ENO+_ ≪ *r*_ENO−_, respectively). This is achieved by the irreversible reactions, which drain and supply metabolites to the ‘super-eno’. Because of the positive feedback between enzyme activity and non-equilibrium of the ‘super-eno’, this regulation topology achieves differential regulation during glycolysis and gluconeogenesis. As we observed in the analysis of the glycolytic and gluconeogenic steady states (Fig. 1), this differential regulation adjusts enzyme levels via transcriptional regulation and suppresses futile cycling at the irreversible reactions.

While regulation of central metabolism efficiently organizes FBP-PEP in a far from equilibrium state during exponential growth, nutrient shifts expose the limitations of this regulatory system. To understand why, we plot FBP against PEP, with both metabolites normalized to their gluconeogenic steady state (Fig. 3A). We indicated several time-points along the dynamics, and the final steady state is shown with a grey symbol. Initially, both FBP and PEP drop close to zero, followed by a very slow joint increase of FBP and PEP over the course of hours (Fig. 3A). This joint increase, rather than a differential increase, is the hallmark of a close-to-equilibrium state.

The slow recovery can be understood from the simulation, which shows that FBP and PEP proceed close to the equilibrium line of Eq. (7), where growth is slow (Fig. 3B). As a guide to the eye, we drew an equilibrium parabola in Fig. 3A along the joint increase, too.

We previously showed that throughout most of the lag phase, higher gluconeogenic flux from increasing levels of gluconeogenic enzymes is almost completely lost to a corresponding increase in futile cycling because increasing FBP activates lower glycolysis, instead of deactivating it (Fig. 2F). The overshoot of FBP in Fig. 3A (data) and Fig. 3B (model) is what finally allows the cell to establish net flux because it is breaking the equilibrium: PEP concentration is high enough to activate upper gluconeogenesis sufficiently to drain FBP via upper gluconeogenesis (see Fig. 2E). Lower FBP then shuts down futile cycling in lower glycolysis/gluconeogenesis (Fig. 2F), pushing FBP and PEP concentrations to a state far from the equilibrium line (see Fig. 3B) and allowing the cell to grow at a faster rate.

The fundamental difference between shifts to gluconeogenesis and glycolysis in *E. coli* is that glycolytic shifts immediately land far from equilibrium (Fig. 3C, triple arrow to white circle), such that cells immediately grow at faster rates, allowing them to express the new enzymes needed to recover quickly. But why does one direction immediately land far from equilibrium, while the other lands close to equilibrium?

### Three trade-offs constrain lag times to glycolysis and gluconeogenesis

The out-of-equilibrium state is caused by net flux going through metabolism. Therefore, we investigated what causes fluxes not to flow in a uniform direction after shifts to glycolysis and gluconeogenesis. In principle, metabolite flux brought to the ‘super-eno’ can exit via two drains: upper gluconeogenesis, activated by PEP, and lower glycolysis, activated by FBP (Fig. 4A). How much flux exits via either drain depends on the current protein abundances and the allosteric regulation. If the allosteric regulation and protein abundances favor the lower drain, then after a switch to glycolysis, FBP builds up, PEP is drained and a net flux is immediately accomplished. In a shift to gluconeogenesis, however, flux that enters central metabolism from the TCA cycle will immediately drain back to the TCA cycle, leading to an in-and-out flux but no net flux. In this situation, FBP and PEP stay in equilibrium and the recovery stalls. If on the other hand, the upper drain was favored over the lower drain, then we would expect the behavior to be reversed and gluconeogenic flux would be immediately accomplished, while the glycolytic recovery would stall.

**Figure 4.**
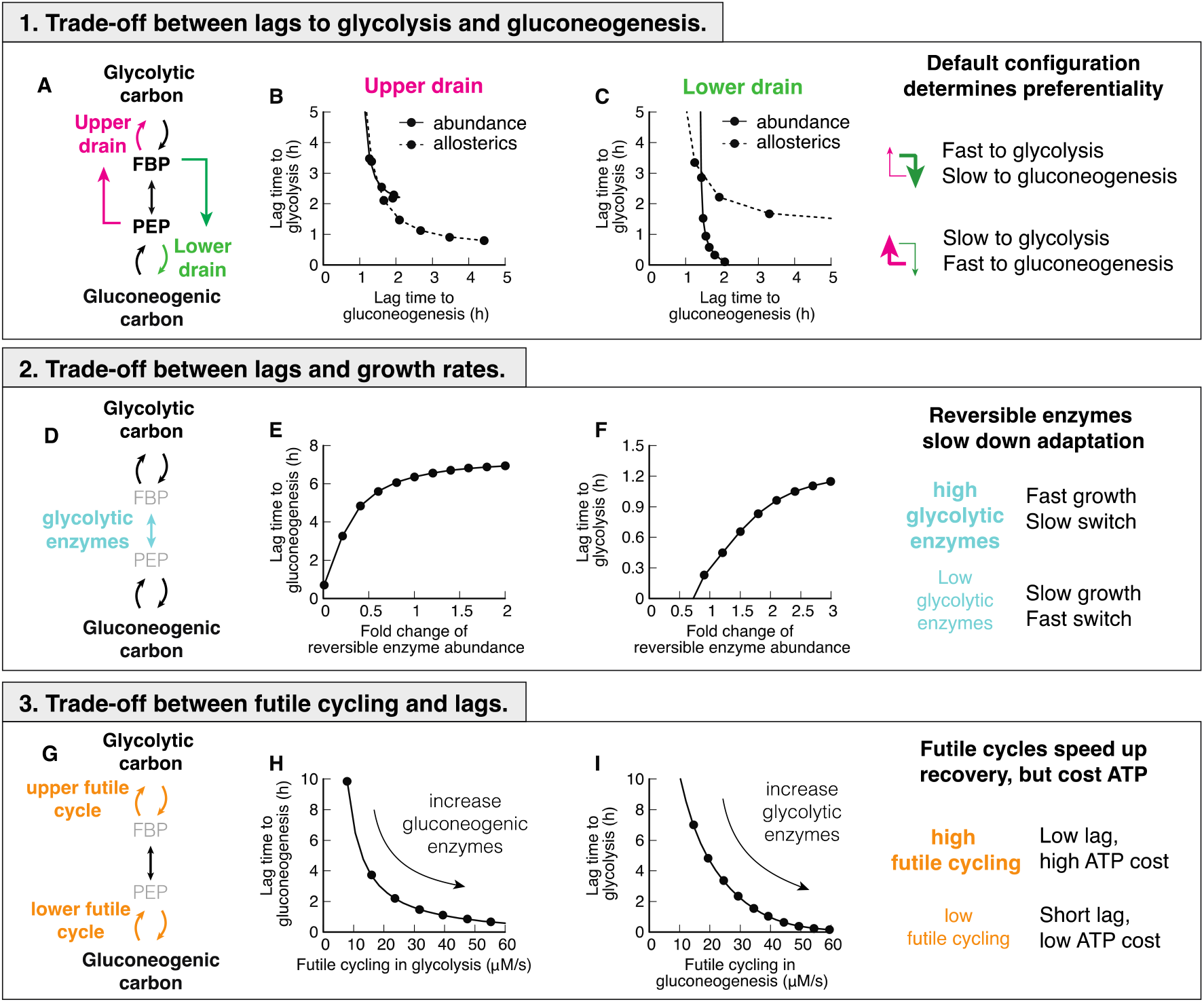
Trade-offs between glycolysis and gluconeogenesis. (**A**) Two drains in central metabolism deplete central metabolites. (**B-C**) Changing abundance ϕ or allosteric regulation strength α in either lower or upper drain leads to a shift of lag times, decreasing lags in one direction at the cost of the other. Chosing strength of the drains such that either top or bottom is stronger, will lead to a fast recovery in on direction, and a slow in the other. (**D**) Reversible enzymes in the central metabolism (coarse-grained here into ‘super-eno’). Abundance of reversible enzymes scale linearly with growth rate [16]. (**E-F**) Decreasing abundance of reversible enzymes decreases lag times. This effect is due to regulatory metabolites being in a far-from-equilibrium state when abundances are low, which allows differential regulation via FBP and PEP. For high abundance, regulation is weak and lag times long. (**G**) There are two futile cycles in central metabolism. (**H-I**) Increasing abundance of enzymes of the opposing direction in pre-shift, e.g. gluconeogenic enzymes in glycolytic growth, increases futile cycling and decreases lag times. Because in futile cycles free energy is dissipated, usually in the form of ATP hydrolysis, futile cycling has an energetic cost.

In the simulation, we are able test the hypothesis that the the upper and lower drains determines the preferential directionality of the central metabolism by varying enzyme abundances and the strength of allosteric interactions in upper and lower drains *in silico*. We let metabolism adapt to gluconeogenic and glycolytic conditions and calculate lag times (Fig. 4B&C). Indeed, we found that a decrease of lag time in one direction led to an increase of lag time in the opposite direction.

Varying the outflow from metabolism is not the only determinant of lag times. The set of reversible enzymes, coarse-grained in our model into the ‘super-eno’, plays another key role because it interconverts the regulatory metabolites FBP and PEP (Fig. 4D). If this conversion is fast, the concentrations of FBP and PEP will be close to their equilibrium relation in Eq. (7), and differential regulation will be impossible. As a result, lag times in both directions increase if we increase the abundance of reversible reactions (Fig. 4E-F). This is a counter-intuitive result, as one would have naïvely expected more enzymes to speed up reactions. But instead, in metabolism more enzymes will collapse the differential regulation and slow down adaptation rates. This trade-off is anavoidable for fast growing cells because the cell needs a sufficient amount of reversible glycolytic enzymes to catalyze metabolic flux.

Finally, lag times depend on the amount of futile cycling, i.e., the circular conversion of metabolites in the upper and lower irreversible reactions (Fig. 4G). Increasing the abundance of gluconeogenic enzymes in glycolytic growth or glycolytic enzymes in gluconeogenic growth increases futile cycling but decreases lag times (Fig. 4I&H). Because futile cycling dissipates ATP, which is not explicitly built into our model, this third trade-off means that organisms can decrease their switching times by sacrificing energetic efficiency.

Are these three trade-offs a fundamental consequence of the regulatory structure or are there parameter combinations that avoid the trade-offs by simultaneously enabling rapid growth and rapid switching without costly futile cycling? To answer this question we performed an extensive scan of model parameters by randomly choosing sets of biochemical parameters and simulating the resulting model. Of those parameter sets we chose those that allowed steady state growth in both glycolytic and gluconeogenic conditions and were able to switch between both states. We plotted the sum of futile cycling in the upper and lower irreversible reactions in the pre-shift conditions against the subsequent lag times for shifts to gluconeogenesis (Fig. 5A) and to glycolysis (Fig. 5B). In addition, we colored individual parameter sets according to the total allosteric regulation, defined as the sum of fold-changes of enzyme activities between glycolysis and gluconeogenesis (black: R < 10^2^, red/green: 10^4^ > R > 10^2^, grey: R > 10^4^). These fold changes are the result of both allosteric and transcriptional variations. We found that metabolism in the majority of randomly generated models is inefficient and dominated by futile cycling; only a minority of models were able to reduce futile cycling in glycolysis and gluconeogenesis. Remarkably, despite probing variations of all possible model parameters, including Michaelis Menten parameters of enzymes and the strengths of allosteric and transcriptional regulation, lag times could not be reduced at-will by the cell. Instead, individual parameter sets with similar allosteric regulation (colors) are bound by a ‘ Pareto frontier’ (solid lines) between futile cycling in preshift conditions and lag times. Points close to the ‘Pareto frontier’ are Pareto-optimal, meaning that any further decrease of either parameter must come at the expense of the other. Overall, stronger allosteric regulation shifted the Pareto frontier but was not able to overcome it. Parameter combinations that led to low futile cycling in either glycolysis or gluconeogenesis showed long lag times in at least one condition (Fig. 5C, ‘black’ and ‘yellow’) compared to the background of all simulated parameter sets (‘grey’). Thus, from this analysis, it seems that organisms with the regulatory architecture of Box 1 cannot overcome long lag times without paying a futile cycling cost during steady state growth.

**Figure 5.**
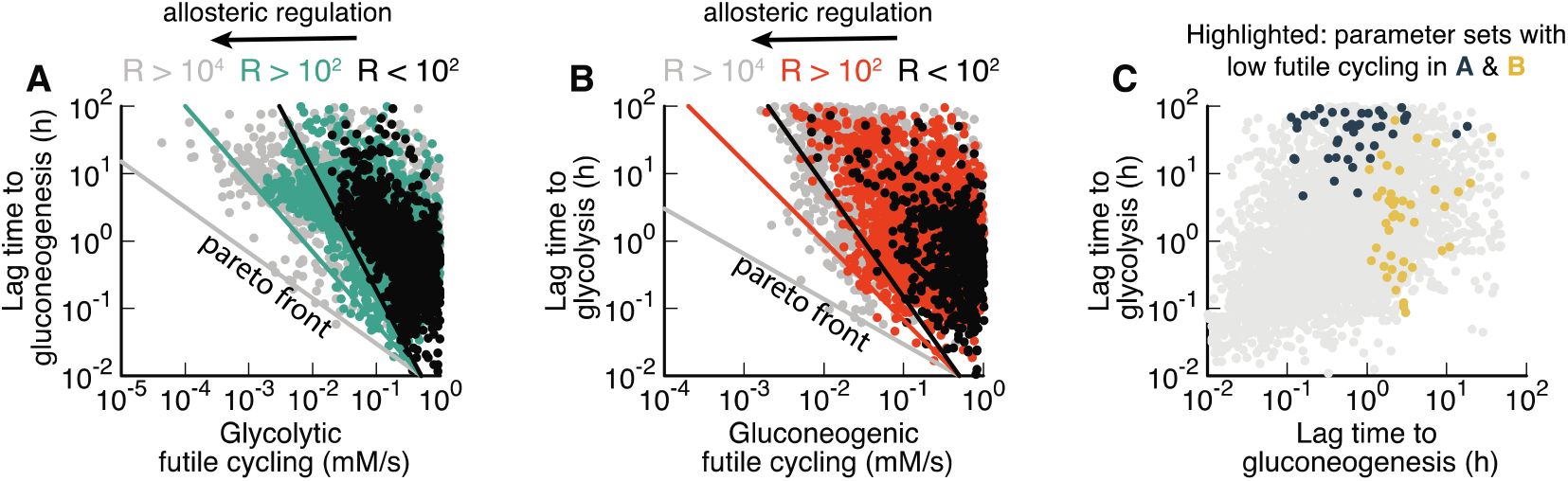
Large-scale parameter scan reveals Pareto optimality between lag times and futile cycling. (**A-B**) Model calculated for randomized sets of protein abundancies, reaction rates, Michaelis constants, allosteric interactions, transcriptional regulation, see SI. Each point corresponds to a parameter set that allows exponential growth on both glycolytic and gluconeogenic carbons, as well switching between both conditions. Data is colored according to the total regulation R, i.e., the sum of fold-changes of enzyme activities between glycolysis and gluconeogenesis, 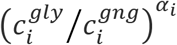, where 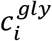 and 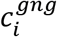 are protein abundances in glycolysis and gluconeogenesis of protein i and α_i_ the strength of the allosteric regulation. For standard E. coli parameters R = 23. R>10^4^ are likely unphysiological. Lines indicate Pareto front and are drawn by hand. (**C**) Parameter sets from panels A&B with low futile cycling highlighted over the background of all parameter sets (grey).

### Gluconeogenesis specialists are constrained by the same trade-offs

Taken together, the results of Fig. 4&5 suggest that microbial cells cannot achieve fast growth, low futile cycling and fast adaptation simultaneously in both glycolysis and gluconeogenesis. Instead, trade-offs between these six extremes constrain the evolutionary optimization of microbial metabolism, such that any optimal solution is on a surface of a multidimensional Pareto frontier, where any improvement in one phenotype will come at the expense of others. To test this hypothesis, we next asked whether a gluconeogenic specialist would indeed be constrained by the same trade-offs as *E. coli* and other glycolytic specialists. For this purpose we chose *P. aeruginosa*, a well-studied gluconeogenesis specialist that has a similar maximal growth rate in minimal medium as *E. coli* (*E. coli* 0.9/h on glucose, *P.aeruginosa* 1.0/h on malate) and grows on a wide variety of substrates.

Strikingly, *P. aeruginosa* grows fast on gluconeogenic substrates that are considered ‘poor’ substrates for *E. coli*, but slow on glycolytic substrates that are considered ‘good’ (Fig. 6A). From our model, we would expect that such a specialization for gluconeogenic substrates would go along with a reversal in lag phases, too. Indeed, switching between glycolytic and gluconeogenic substrates, *P. aeruginosa* exhibits a mirrored pattern of lag phases compared to *E. coli* (compare Fig. 6B to 6C), with a long multi-hour lag phase when switched to glycolysis.

**Figure 6.**
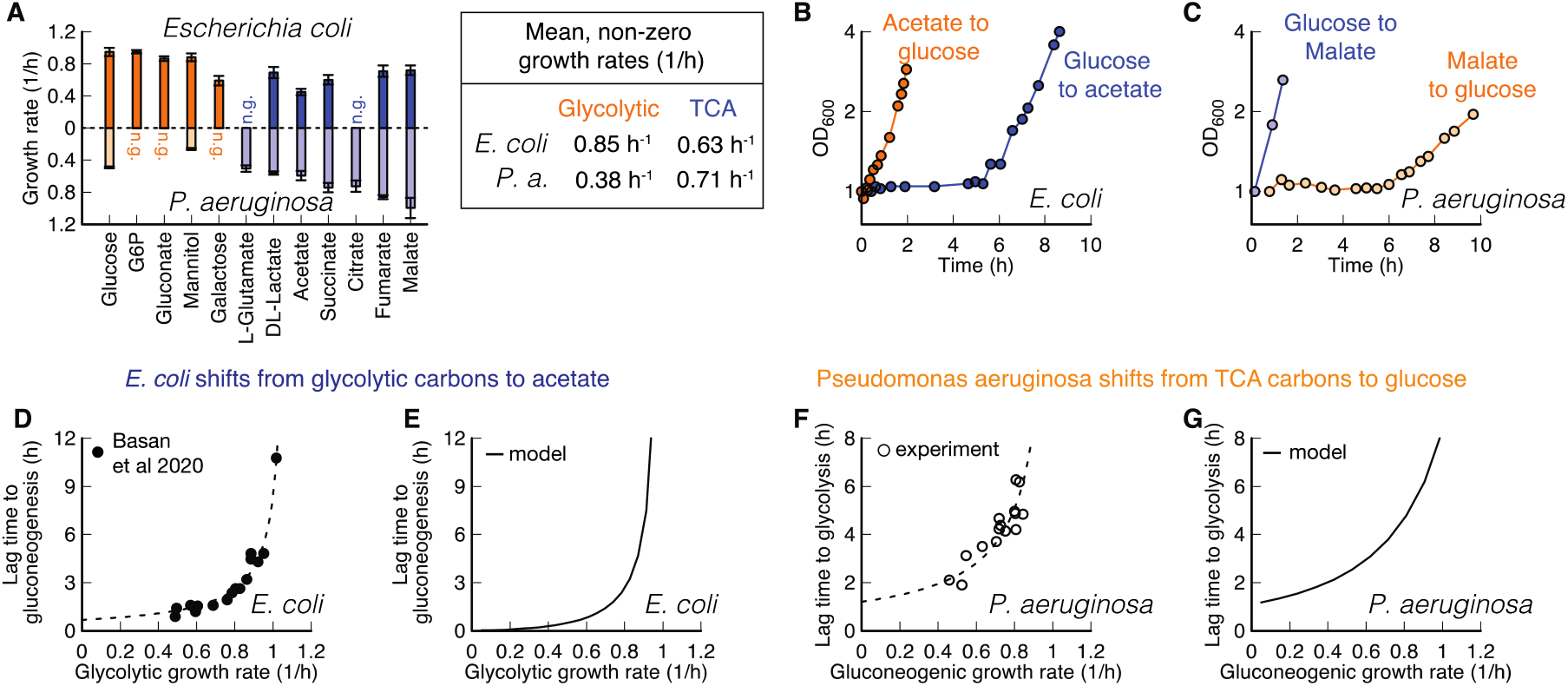
Comparison of *E. coli* and *P. aeruginosa* during growth and shifts. (**A**) Growth rates on glycolytic carbons (orange) are faster for *E. coli* than on gluconeogenic carbons (blue). For Pseudomonas, this dependence is reversed. No growth indicated with “n.g”. Data is average of n=3. Error bar is standard deviation. (**B-C**) Shifts for *E. coli* and *P. aeruginosa* between glycolytic and gluconeogenic carbon substrates. The preferential order of P. aeruginosa is reversed in comparison to *E. coli* (**D**) *E. coli* shows an increase of lag times to gluconeogenesis with increasing pre-shift growth rate. Lag times diverge around growth rate 1.1/h. Each point is an individual experiment. (**E**) The model predicts diverging growth rates without further fitting, based on the growth rate dependent expression levels of glycolytic and gluconeogenic enzymes (Fig. 2E-F). (**F**) P. aeruginosa shows a strikingly similar growth rate to lag time dependence as *E. coli*, when switched to glycolysis, with lag times diverging around 1.0/h. Each point is an individual experiment. (**G**) The model can recapitulate observed *P. aeruginosa* lag times if pre-shift glycolytic enzymes are decreased as a function of pre-shift growth rate.

To investigate if both *E. coli* and *P. aeruginosa* are constrained by the same trade-offs, we investigated the effect of pre-shift growth rate, which according to Fig. 4 should have a negative effect on growth rates. For *E. coli* it is known that shifts from glycolysis to gluconeogenesis depend on the pre-shift growth rate (Fig. 6D, data: (Basan et al., 2020)), which we can capture in our model if we take FBP-dependent transcriptional regulation into account (Fig. 6E). We tested the corresponding lag times for *P. aeruginosa* by varying gluconeogenic substrates and found a similar dependency in shifts to glycolytic substrates (Fig. 6F&G). Hence as expected from the model, these findings show that *P. aeruginosa* is constrained by the same trade-offs as *E. coli*.

To decipher whether *P. aeruginosa* lag times are constrained on a molecular level by the same inability to break the equilibrium after nutrient shifts, we investigated metabolite concentration dynamics in central metabolism. Because *P. aeruginosa_uses* the ED pathway for hexose catabolism (Wang et al., 1959; Vicente and Cánovas, 1973), we needed to adapt our model slightly. The irreversible reactions in the ED pathway convert gluconate-6-phosphate to glyceraldehyde 3-phosphate (GAP) and pyruvate. In the reversible chain of reactions, the first metabolite in glycolysis is thus GAP rather than FBP. Because GAP is difficult to quantify in mass spec-based metabolomics, we used the closely related compound dihydroxyacetone phosphate (DHAP) as a proxy. DHAP is in chemical equilibrium with GAP via a single fast and reversible isomerase (Nikel et al., 2015).

Analogous to Fig. 3, we plot the dynamics of DHAP versus PEP, normalized to their glycolytic steady state values, for both shifts (Fig. 7A&B). The dynamics starts and ends at their respective steady states (grey symbols and dashed lines) and follows the direction of the indicated arrow. In the chemical equilibrium, DHAP depends linearly on PEP, 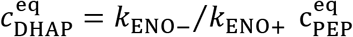, analogous to Eq. (7), but without the square because of the 1-to-1 stochiometry between DHAP and PEP. This equilibrium is indicated with a red line. During the long lag time of *P. aeruginosa* in a shift from malate to glucose, we see that initially both DHAP and PEP collapse, followed by a slow increase along the equilibrium line (Fig. 7A). Thus, despite substantial amounts of metabolites being built-up, ‘super-eno’ remains close to equilibrium. Only after 5.6 h, when the DHAP-PEP dynamics deviates from the line, the equilibrium is broken and net flux can be achieved.

**Figure 7.**
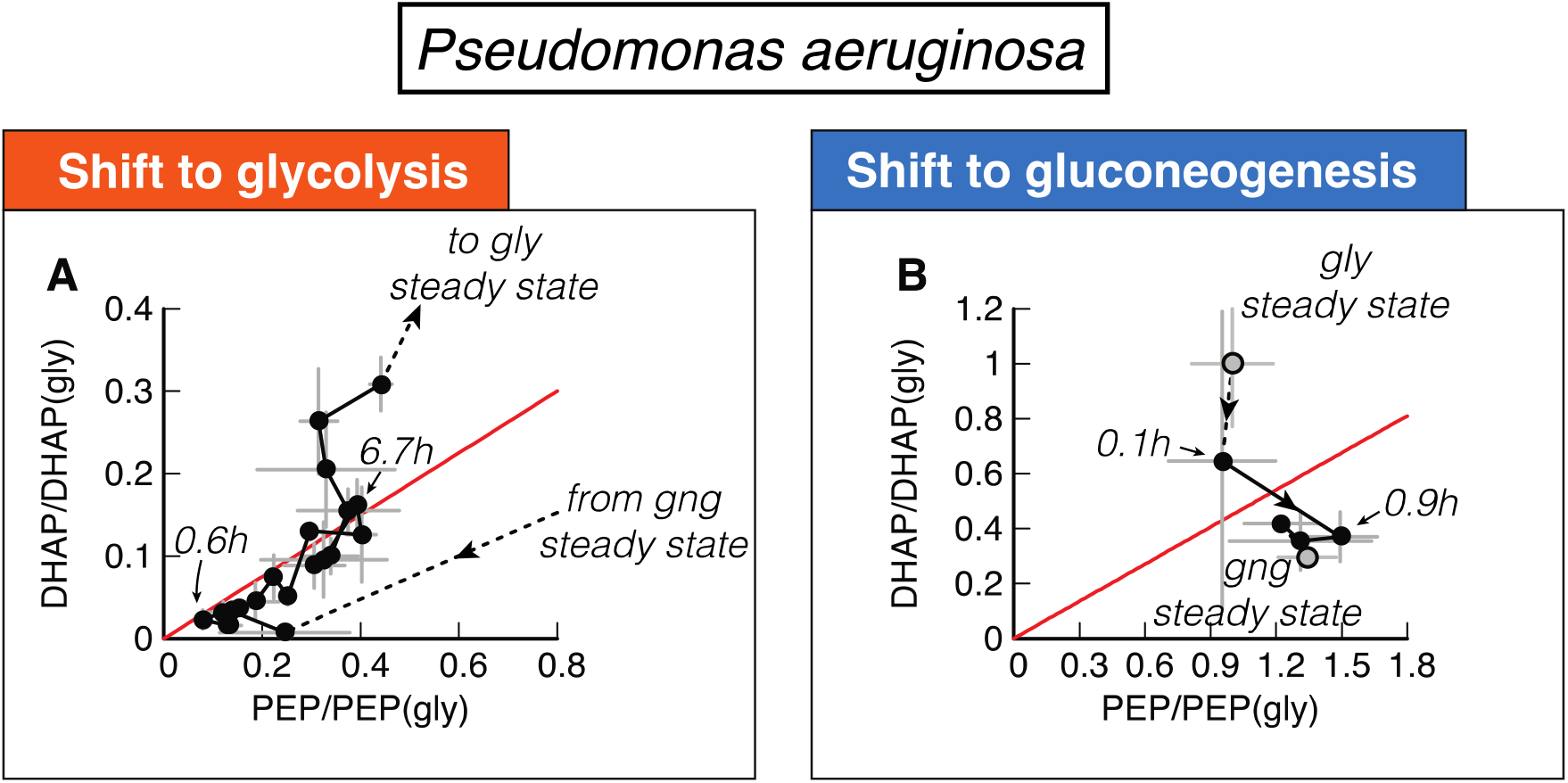
Metabolite dynamics of *P. aeruginosa* during shifts from malate to glucose and vice-versa. (**A**) DHAP and PEP during shift from malate (‘gng’) to glucose (‘gly’), normalized to the final glycolytic steady state. Recovery follows a direct proportionality, indicating that central metabolism is close to equilibrium (red line) during the recovery. (**B**) DHAP and PEP reach the final steady state (‘gng’) without creeping along the equilibrium line. Data is the average of n=3, error bars show standard deviation.

In the reverse shift from glucose to malate, *P. aeruginosa*, in constrast, can immediately establish a non-equilibrium and grow. Thus not only is the asymmetry in lag times reversed compared to *E. coli*, it is also caused by the same inability to break the equilibrium and establish net flux in central metabolism.

But, microbes do not have to be optimized for either direction. One such case is *P. putida* with moderate lag times of about 1 to 2 h in both directions and only a slight preference for gluconeogenic susbtrates (EV Fig. 2). According to the model, such a generalist strategy can also be a Pareto-optimal solution of the biochemical trade-offs of Fig. 4-5, but it must come at the expense of no fast recovery (Fig. 4A-C) and reduced growth because of the trade-offs with reversible enzymes (Fig. 4D-F) and futile cycling (Fig. 4G-I). This is indeed the case for *P. putida*. Lag times are in the fast direction are twice as long compared to *P. aeruginosa* and the growth rate is about 20% slower (Fig. S8).

## Discussion

In this work, we presented a coarse-grained kinetic model of central carbon metabolism, combining key allosteric and transcriptional regulation, as well as biomass production, enzyme synthesis, and growth. This model elucidates the remarkable capacity of central carbon metabolism to self-organize in response to substrate availability and flux requirements. During exponential growth, regulatory metabolites adjust to far-from-equilibrium steady states, providing the cell with an elegant mechanism to sense the required directionality of the flux. But the model reveals a key limitation of this flux sensing. Because after a nutrient shift the concentration of the metabolites collapses to its equilibrium, the cell becomes ‘blind’ to the direction that the flux is supposed to flow through the system. By implementing a preferred direction, the cell can partially overcome lag times in one direction at the cost of increasing lag times in the opposite direction. In addition, two more trade-offs constrain the ability to simultaneously decrease both lag times, because it impacts growth rate and the level of futile cycling during growth.

Microbial species can maximize their proliferation only up to the Pareto-frontier spanned by these trade-offs, which can lead to evolution of substrate specialization. We validated this key model prediction in different bacterial species. In *P. aeruginosa* we showed a reversal of substrate preference as compared to *E. coli*, which coincided with a complete reversal of the phenomenology of lag phases and metabolite dynamics. In *P. putida* we found a generalist strategy with moderate lag times in both direction.

One of the results from our model is that lag times could be substantially reduced by allowing futile cycling, e.g., by expressing irreversible enzymes for both directions at all times. The energetic cost of such a wasteful strategy would be relatively low. Because energy production pathways only constitute a relatively small fraction (around 20% (Basan et al., 2015c)) of the total cellular proteome and nutrient uptake even smaller (around 1% for glucose uptake (Schmidt et al., 2016)), the cell could compensate ATP dissipated in futile cycling by increasing nutrient uptake and ATP production at a relatively low proteome cost. However, experimentally it appears that *E. coli* chooses to keep futile cycling in check by transcriptionally regulating irreversible enzymes. We thus hypothesize that the cost of futile cycling must be considered in conditions where the energy budget is much more limited, such as growth shifts and during starvation. In fact, it has been recently shown that the energy budget of the cell is around 100-fold smaller during carbon starvation and that energy dissipation can increase death rates several-fold (Schink et al., 2019). Therefore, even levels of futile cycling that are modest during steady state growth should severely affect survival of cells in these conditions

Our findings indicate that the identified trade-offs are inherent properties of central carbon metabolism, at least given the existing allosteric and transcriptional regulation. But could different regulation overcome this limitation? In theory, the cell could use a direct input signal from the carbon substrate to allosterically inhibit or even degrade undesired metabolic enzymes. This would uncouple enzyme abundances and activies in pre- and post-shift growth and circumvent the trade-offs. But with dozens of glycolytic and gluconeogenic substrates, this would result in a much higher degree of regulatory complexity, quickly exceeding the regulatory signal capacity that microbes with their small genomes could sense and integrate. In addition, any wrong decision to degrade or inhibit metabolic enzymes, for example when combinations of nutrients are present or when supply is only briefly inhibited, would drastically impair growth. Thus the regulatory network that microbes use might not be maximizing growth, but at least it is robust and prevents misregulation.

Another reason why no such regulation has evolved could be related to the observation that the regulation of upper and lower glycolysis/gluconeogenesis and directionality of flux are performed by the metabolite concentrations of FBP and PEP, which are cut off from the rest of metabolism by irreversible reactions. We propose that the logic for this regulatory architecture is product inhibition, which ensures that this essential part of central carbon metabolism is adequately supplied with metabolites, but also ensures that uncontrolled and potentially toxic accumulation of metabolites does not occur. In fact, because the reactions of upper and lower glycolysis are effectively irreversible, even a slight misbalance in flux between these enzymes and biomass demand would result in uncontrolled accumulation of metabolites and, in the absence of a cellular overflow mechanism, these metabolites would quickly reach toxic concentrations, e.g., via their osmotic activities. As demonstrated by the simulation, the existing regulation of central metabolism successfully resolves this problem.

The regulatory architecture of central metabolism accomplishes efficient regulation of fluxes and metabolite pools in response to diverse external conditions while avoiding toxic accumulation of internal metabolites and integrating multiple conflicting signals with only two regulatory nodes. Central metabolism is a remarkable example of self-organization of regulatory networks in biology. It provides an elegant solution to the complex, obligatory problem, posed by the biochemistry of central carbon metabolism. All organisms that need to switch between glycolytic and gluconeogenic flux modes face this problem, and we argue that this explains the striking degree of conservation of the phenomenology of shifts between glycolytic and gluconeogenic conditions that we found in different microbial species, ranging from *E. coli, B. subtilis*, and even wild-type strains of the lower eukaryote *S. cerevisiae* to the reversed phenotypes in *P. aeruginosa*. Conversely, we believe that the quantitative phenotypes exhibited by microbes in such idealized growth shift experiments in the lab can reveal much about their natural environments, ecology and evolutionary origin.

## Code and data availability

The datasets and computer code produced in this study are available in the following databases:

- The MATLAB implementation of the model and the data of all figures are available on github: https://github.com/Severin-schink/Glycolysis-gluconeogenesis-switches
- Metabolomics data is available on Metabolights: www.ebi.ac.uk/metabolights/MTBLS3887

## Conflict of interest statement

The authors declare that they have no conflict of interest.

## Acknowledgments

We thank Terence Hwa for many helpful discussions and always pointing out unresolved questions in microbial physiology. We thank Mark Polk for proofreading the manuscript. This project was financed by MIRA grant (5R35GM137895) and an HMS Junior Faculty Armenise grant via MB and HFSP Long-term fellowship (LT000597/2018) via SJS. EA was supported by HCRP and the Program for Research in Science and Engineering (PRISE) of Harvard.

## Author contributions

SJS, DC, AM, TF, US and MB contributed to the design of the project and writing the manuscript. SJS, DC, EA and MB performed modelling. AM and MB performed growth experiments. VB, TF and GAB performed metabolomics measurments.

## Materials and methods

### Bacterial cultures

Strains used in this paper are wild-type *Escherichia coli* K-12 NCM3722 (Soupene et al., 2003), *Pseudomonas aeruginosa* PAO1 (Stover et al., 2000) and *Pseudomonas putida* NIST0129. The culture medium used in this study is N^-^C^-^ minimal medium (Csonka et al., 1994), containing K_2_SO_4_ (1 g), K_2_HPO_4_·3H_2_O (17.7 g), KH_2_PO_4_ (4.7 g), MgSO_4_·7H_2_O (0.1 g) and NaCl (2.5 g) per liter. The medium was supplemented with 20 mM NH_4_Cl, as the nitrogen source, and either of the following carbon sources: 20 mM Glucose-6-phosphate, 20 mM gluconate, 0.2 % glucose, 20 mM succinate, 20 mM acetate, 20 mM citrate, 20 mM malate or 20 mM fumerate.

Growth was then carried out at 37 °C in a water bath shaker at 200 rpm, in silicate glass tubes (Fisher Scientific) closed with plastic caps (Kim Kap). Cultures spent at least 10 doublings in exponential growth in pre-shift medium. For growth shifts, cultured were transferred to a filter paper and washed twice with pre-warmed post-shift medium. Cells were resuspended from the filter paper in post-shift medium and subsequently diluted to an OD of about 0.05.

### Preparation of metabolite samples

Each metabolite sample was filtered, and the filter was immediately plunged in 4 ml ice cold Methanol (40 %)+Acetonitrile (40 %)+water (20 %) and kept in 50 ml tube. Bacteria were washed off from the filter by pipetting, and the solution was transferred to 15 ml tube. Original 50 ml tube was further washed with 4 ml of ice cold Methanol+Acetonitrile+Water mix and added to respective 15 ml tube (total 8 ml). Each sample was dried by speed vac, and dried extracts were sent for Mass spec analysis.

### Quantification of intracellular metabolite levels

The dried metabolite extracts were resuspended in 150 μL MilliQ water, centrifuged at 4 °C, 10,000 rpm for 10 min, and 100 μL precipitate-free supernatant was transferred to a master 96-well plate. 25 μL of the master plate were transferred to a 96-well plate for acquisition, of which 10 μL were injected into a Waters Acquity ultraperformance liquid chromatography (UPLC) system (Waters) with a Waters Acquity T3 column coupled to a Thermo TSQ Quantum Ultra triple quadrupole instrument (Thermo Fisher Scientific) as described previously (Buescher et al., 2010). Compound separation was achieved using a gradient of two mobile phases: A, 10 mM tributylamine (ion-pairing agent), 15 mM acetate and 5% (*v*/*v*) methanol in water; and B, 2-propanol. Data was acquired in negative ionization mode using previously published MRM settings (Buescher et al., 2010). Peak integration was performed using an in-house software based on MatLab. A dilution series of standards was used to calculate the concentrations of metabolites in the samples. The final intracellular concentration was calculated from the sample concentration and the extracted intracellular volume.

### Theoretical modelling

The integrated minmal model of metabolism and growth was implemented in MATLAB using the SimBiology toolbox and is described in detail in the Supporting Information.

## Expanded View figures

**EV Figure 1.**
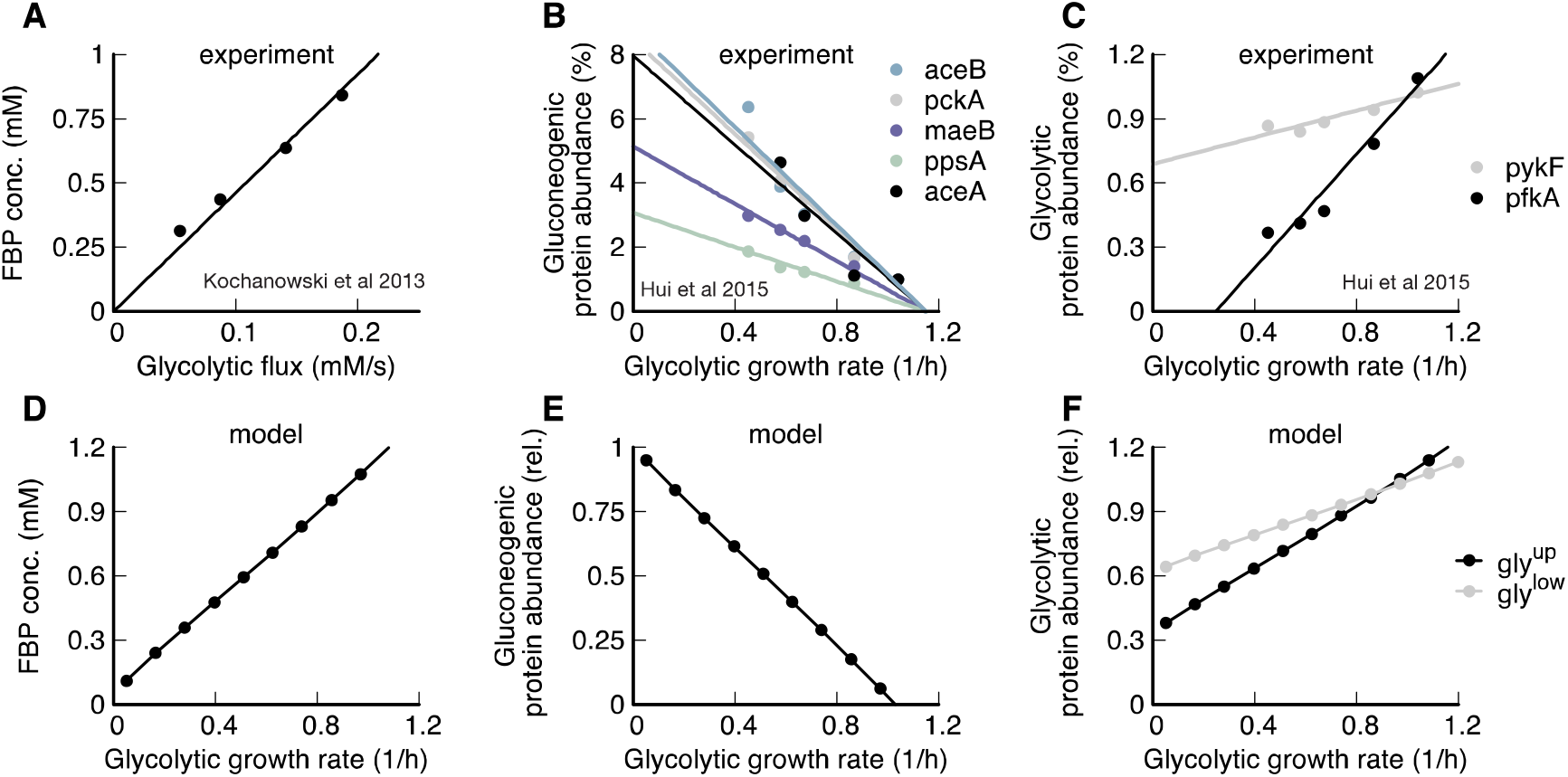
Metabolic state depends on growth rate. (**A**) During glycolytic growth, FBP linearly increases with growth rate. Data: Ref. (Kochanowski et al., 2013b). (**B**) Gluconeogenic enzymes decrease linearly with glycolytic growth rate. Data: (Hui et al., 2015). (**C**) Glycolytic enzymes increase linearly with glycolytic growth rate. Data: Ref. (Hui et al., 2015). (**D-F**) Simulation results recapitulate experimental evidence.

**EV Figure 2.**
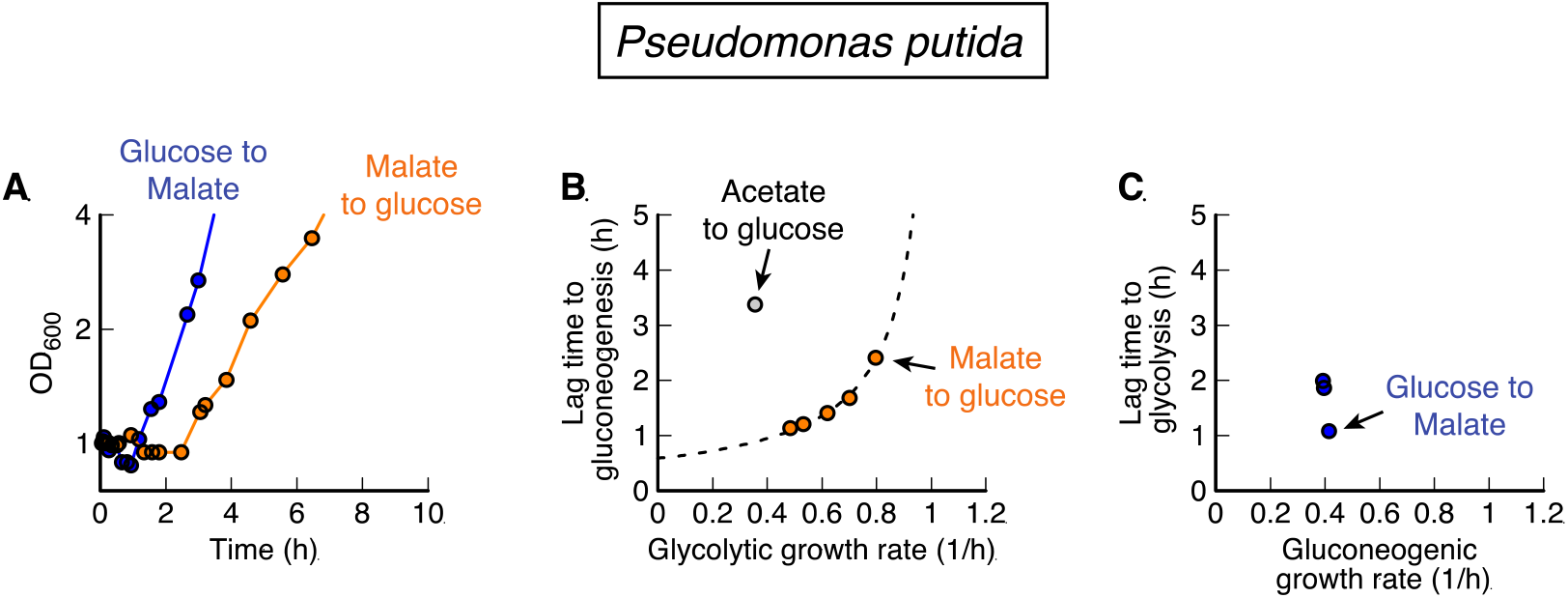
Pseudomonas putida shifts between glycolytic and gluconeogenic carbon substrates. (**A**) For shifts between glucose and malate, and vice-versa, P. putida shows moderate lag times in both directions. (**B**) Lag times depend on pre-shift growth rate for glycolysis to gluconeogenesis, with acetate to glucose being a clear outlier. (**C**) Lag times for gluconeogenic to glycolytic shifts are of the same magnitude as for the reverse direction. Compared to P. aeruginosa, P. putida shows longer lags from glucose to malate (0.55h compared to 1.1h) and slower growth on the fastest carbon (malate) (1.0/h compared to 0.8/h)

## Notes

### Competing Interest Statement

The authors have declared no competing interest.

