## Supplemental Information for "Glycolysis/gluconeogenesis specialization in microbes is driven by biochemical constraints of flux sensing"

### Appendix of
### Substrate specialization in microbes is caused by biochemical constraints of dynamic flux sensing

Severin Josef Schink[1,*], Dimitris Christodoulou[1,2,*], Avik Mukherjee[1,3], Edward Athaide[1], Viktoria Brunner[2], Tobias Fuhrer[2], Gary Andrew Bradshaw[4], Uwe Sauer[2, #] and Markus Basan[1, #]

* Equal contribution

[1] Systems Biology Department, Harvard Medical School, 200 Longwood Ave, Boston, MA 02115, USA

[2] Institute of Molecular Systems Biology, ETH Zurich, Zurich 8093, Switzerland

[3] Applied Mathematics Department, Harvard College, Cambridge, MA 02138, USA

[4] Laboratory of Systems Pharmacology, Harvard Program in Therapeutic Science, Harvard Medical School, 200 Longwood Ave, Boston, MA 02115, USA

### 1. Introduction

In the following notes we describe the coarse-grained model of central carbon metabolism outlined in Box 1 and used throughout this paper. We start with metabolic reactions. Next, we derive how these metabolites lead to biomass growth and combine it with proteome allocation to derive changes in enzyme abundancies. After having detailed the metabolic model, we will outline our approach to solve it using a combination of analytical and numerical methods. Finally, we narrow down the parameter space and describe how we chose the parameter set used in this paper.

### 2. Coarse-grained model of central carbon metabolism

The design idea behind the model in this paper is to coarse grain the complexity of central carbon metabolism to its two defining features during growth transitions: the irreversible reactions of Fructose 6-phosphate (F6P) to Fructose-1,6-bis-phosphate (FBP) and Phosphoenolpyruvate (PEP) to Tricarboxylic acid cycle carbons (TCA). Each of these two reactions is catalyzed in the forward and backward direction by distinct enzymes. In contrast, most other enzymes in central metabolism are reversible, meaning that forward and backward reactions are catalyzed by the same enzyme.

#### 2.1. Differential equations of the model

The model is governed by a series of differential equations involving the concentrations of metabolites and abundancies of enzymes.

#### 2.2. Glycolytic in- and outflux

Our model coarse-grains glycolytic carbons, such as F6P or glucose, into a single metabolite "GLY", that is taken up, depending on the availability of a glycolytic carbon

$$r_{\emptyset\rightarrow\mathrm{GLY}} = \mathrm{V}_{\mathrm{max}}^{\mathrm{gly,up}}\, \mathbb{1}_{\mathrm{gly}} \tag{S1}$$

for uptake, with $V_{\mathrm{max}}^{\mathrm{gly,up}}$ being the maximum possible uptake rate of glucose and $\mathbb{1}_{\mathrm{gly}}$ being an indicator function for the presence of glycolytic carbon. In addition, glycolytic carbons can also be drained in our model, to prevent build-up of metabolites after nutrient shifts,

$$r_{\mathrm{GLY}\to\emptyset} = V_{\max}^{\mathrm{gly,drain}}\,[\mathrm{GLY}] \tag{S2}$$

for drainage, with $V_{\max}^{\mathrm{gly,drain}}$ being a proportionality constant and [GLY] here referring to the concentration of glycolytic carbons.

##### 2.3. Upper irreversible reactions

The pair of reactions between glycolytic carbons and FBP has rates that are controlled by allosteric regulation by PEP. In the SI, we call the irreversible reactions by the names of their respective enzymes, i.e. pfk and fbp for upper glycolysis and gluconeogenesis. The reaction rates of these enzymes are calculated as

$$r_{\mathrm{GLY}\to\mathrm{FBP}} = k_{\mathrm{cat}}^{\mathrm{pfk}}\,\phi_{\mathrm{pfk}}\left(\frac{[\mathrm{PEP}]}{[\mathrm{PEP}^*]}\right)^{-\alpha^{\mathrm{pfk}}}\frac{[\mathrm{GLY}]}{K_{\mathrm{M}}^{\mathrm{pfk}}+[\mathrm{GLY}]} \tag{S3}$$

and

$$r_{\mathrm{FBP}\to\mathrm{GLY}} = k_{\mathrm{cat}}^{\mathrm{fbp}}\,\phi_{\mathrm{fbp}}\left(\frac{[\mathrm{PEP}]}{[\mathrm{PEP}^*]}\right)^{\alpha^{\mathrm{fbp}}}\frac{[\mathrm{FBP}]}{K_{\mathrm{M}}^{\mathrm{fbp}}+[\mathrm{FBP}]} \tag{S4}$$

where $\alpha^{\mathrm{pfk}}$ and $\alpha^{\mathrm{fbp}}$ are allosteric regulation constants, $k_{\mathrm{cat}}$ is the turnover rate and $\phi$ is the protein abundance of the respective enzyme. This formulation of reaction rates is a modified Michaelis-Menten kinetics, with the allosteric regulation being modeled by a Hill function in the regime of metabolite concentrations being below the half-saturation point of the Hill function.

##### 2.4. Reaction rates of reversible super-eno reaction

The reactions between FBP and PEP is modeled using substrate-limited kinetic laws, so their rates are

$$r_{\mathrm{FBP}\to 2\mathrm{PEP}} = k_{\mathrm{cat}}^{\mathrm{f}}\phi_{\mathrm{eno}}[\mathrm{FBP}] \tag{S5}$$

and

$$r_{2\mathrm{PEP}\to\mathrm{FBP}} = k_{\mathrm{cat}}^{\mathrm{r}}\phi_{\mathrm{eno}}[\mathrm{PEP}]^2 \tag{S6}$$

for the forward and reverse reactions, where $k_{\mathrm{cat}}^{\mathrm{f}}$ and $k_{\mathrm{cat}}^{\mathrm{r}}$ are the catalytic rates of the super eno 'enzyme'. The ratio of rates is set by the dissociation constant $K_{\mathrm{D}} = \frac{k_{\mathrm{cat}}^{\mathrm{r}}}{k_{\mathrm{cat}}^{\mathrm{f}}}$ and cannot be freely chosen by the cell. The abundance of the super-eno enzyme however, $\phi_{\mathrm{eno}}$, can be chosen, and will affect both forward and backward rates simultaneously.

#### 2.5. Lower irreversible reaction

In the lower part of central metabolism, a set of reactions convert PEP into pyruvate and other gluconeogenic carbons and vice-versa. We name the reactions after the two major enzymes that produce pyruvate and PEP, pyk and pck, respectively. We coarse grain reactions in either direction into a single reaction step, and model the dynamics as

$$r_{\mathrm{PEP}\to\mathrm{GNG}} = k_{\mathrm{cat}}^{\mathrm{pyk}} \phi_{\mathrm{pyk}} \left(\frac{[\mathrm{FBP}]}{[\mathrm{FBP}^*]}\right)^{\alpha^{\mathrm{pyk}}} \frac{[\mathrm{PEP}]}{K_{\mathrm{M}}^{\mathrm{pyk}} + [\mathrm{PEP}]} \tag{S7}$$

where $\alpha^{\mathrm{pyk}}$ is a constant for allosteric regulation, $k_{\mathrm{cat}}^{\mathrm{pyk}}$ the catalytic rate and $\phi_{\mathrm{pyk}}$ the enzyme abundance. $[\mathrm{FBP}^*]$ is the FBP concentration in a reference steady state. The rate of conversion of gluconeogenic carbons to PEP is

$$r_{\mathrm{GNG}\to\mathrm{PEP}} = k_{\mathrm{cat}}^{\mathrm{pck}} \phi_{\mathrm{pck}} \left(\frac{[\mathrm{FBP}]}{[\mathrm{FBP}^*]}\right)^{-\alpha^{\mathrm{pck}}} \frac{[\mathrm{GNG}]}{K_{\mathrm{M}}^{\mathrm{pck}} + [\mathrm{GNG}]} \tag{S8}$$

with [GNG] being the concentration of gluconeogenic carbons, e.g., acetate or TCA carbons.

#### 2.6. Gluconeogenic in- and outflux

Lastly, the rates of drainage of gluconeogenic carbon, which coarse-grains pyruvate, acetate and GNG carbons, are given by

$$r_{\mathrm{GNG}\to\emptyset} = V_{\mathrm{max}}^{\mathrm{gng,drain}} [\mathrm{GNG}] \tag{S9}$$

and

$$\mathrm{r}_{\emptyset\to\mathrm{GNG}} = \mathrm{V}_{\mathrm{max}}^{\mathrm{gng,up}} \mathbb{1}_{\mathrm{gng}} \tag{S10}$$

in the same form as uptake and drainage of glycolytic carbon. Here, $\mathbb{1}_{\mathrm{gng}}$ is an indicator function that toggles from zero to one in the presence of a gluconeogenic carbon (GNG).

#### 2.7. Biomass production

Finally, we implement a reaction as a proxy for the formation of biomass from glycolytic carbons, PEP, and gluconeogenic carbon, with the reaction rate

$$r_{\mathrm{BM}} = k_{\mathrm{BM}} \frac{[\mathrm{GLY}][\mathrm{PEP}][\mathrm{GNG}]}{[\mathrm{GLY}^*][\mathrm{PEP}^*][\mathrm{GNG}^*]} \tag{S11}$$

with $k_{\mathrm{BM}}$ being a proportionality constant that converts metabolite flux to biomass. In Section 3 we discuss how to derive $k_{\mathrm{BM}}$ and other biological parameters.

#### 2.8. Metabolite dynamics

Using the above reaction kinetics, we can construct the differential equations governing metabolite concentrations over time. We omit dilution terms that take into account the increase of volume as the cell grows, because cell growth (~ 45 minutes) is much slower than metabolite turnover (few minutes). In addition to all the variables mentioned above, $\beta_{\mathrm{GLY}}$, $\beta_{\mathrm{PEP}}$, and $\beta_{\mathrm{GNG}}$ are the proportions (by mass) of biomass synthesized from glycolytic carbon, PEP, and gluconeogenic carbons, respectively, see Section 3 for details.

We derive the glycolytic carbon change in concentration.

$$\frac{\mathrm{d}[\mathrm{GLY}]}{\mathrm{d}t} = r_{\emptyset\to\mathrm{GLY}} - r_{\mathrm{GLY}\to\emptyset} - r_{\mathrm{GLY}\to\mathrm{FBP}} + r_{\mathrm{FBP}\to\mathrm{GLY}} - \beta_{\mathrm{GLY}} r_{\mathrm{BM}} \tag{S12}$$

FBP change in concentration.

$$\frac{\mathrm{d}[\mathrm{FBP}]}{\mathrm{d}t} = r_{\mathrm{GLY}\to\mathrm{FBP}} - r_{\mathrm{FBP}\to\mathrm{GLY}} - r_{\mathrm{FBP}\to 2\mathrm{PEP}} + r_{2\mathrm{PEP}\to\mathrm{FBP}} \tag{S13}$$

PEP change in concentration

$$\frac{\mathrm{d}[\mathrm{PEP}]}{\mathrm{d}t} = r_{\mathrm{GNG}\to\mathrm{PEP}} - r_{\mathrm{PEP}\to\mathrm{GNG}} + 2\, r_{\mathrm{FBP}\to 2\mathrm{PEP}} - 2\, r_{2\mathrm{PEP}\to\mathrm{FBP}} - \beta_{\mathrm{PEP}} r_{\mathrm{BM}} \tag{S14}$$

Lastly, GNG carbon change in concentration.

$$\frac{\mathrm{d[GNG]}}{\mathrm{d}t} = r_{\emptyset\rightarrow\mathrm{GNG}} - r_{\mathrm{GNG}\rightarrow\emptyset} - r_{\mathrm{GNG}\rightarrow\mathrm{PEP}} + r_{\mathrm{PEP}\rightarrow\mathrm{GNG}} - \beta_{\mathrm{GNG}} r_{\mathrm{BM}} \quad \text{(S15)}$$

#### 2.9. Protein dynamics

In addition to the reactions, the model also accounts for transcriptional regulation of the enzymes necessary for metabolism, governed by differential equations that take into account synthesis and dilution by growth. Generally, the rate of change in activity is given by

$$\frac{\mathrm{d}\phi_j}{\mathrm{d}t} = \mu\left(\chi_j(t) - \phi_j(t)\right), \quad \text{(S16)}$$

where $\chi_j(t)$ is the regulation function, i.e. the fraction of protein synthesis that is protein $j$, and $\phi_j(t)$ is the abundance, i.e. the fraction of the total protein that is protein $j$. In steady state the proteome abundance will equal the regulation function $\phi_j^* = \chi_j^*$. We chose the regulation function based on experimental evidence presented in Fig. 2, that 1) glycolytic and gluconeogenic irreversible enzyme abundance scales linear plus offset and offset minus linear, respectively, and 2) FBP scales linearly with growth rate. As functional form of the regulation function we pick $\chi_j(t) = \phi_j^*\left(\left(1 + x_j\right) - x_j \frac{[\mathrm{FBP}](t)}{[\mathrm{FBP}^*]}\right)$, such that in glycolytic steady state, enzyme abundance automatically equals $\phi_j^*$. The factor $x_j$ can be freely chosen to adjust the slope of the dependence without changing the steady state abundance. The resulting dynamic equation of the gluconeogenic enzyme abundance is given as

$$\frac{\mathrm{d}\phi_j}{\mathrm{d}t} = \mu\left(\phi_j^*\left(\left(1 + x_j\right) - x_j \frac{[\mathrm{FBP}]}{[\mathrm{FBP}^*]}\right) - \phi_j(t)\right). \quad \text{(S17)}$$

For glycolytic enzymes, expression increases with FBP concentration, thus we chose $\chi_i(t) = \phi_i^* \left( x_i + (1 - x_i) \frac{[\mathrm{FBP}](t)}{[\mathrm{FBP}^*]} \right)$, such that enzyme abundance will increase with FBP concentration and glycolytic growth rate. The dynamics equation of the glycolytic enzyme abundance then reads,

$$\frac{\mathrm{d}\phi_i}{\mathrm{d}t} = \mu \left( \phi_i^* \left( x_i + (1 - x_i) \frac{[\mathrm{FBP}]}{[\mathrm{FBP}^*]} \right) - \phi_i(t) \right). \tag{S18}$$

#### 2.10. Biomass growth, growth rate and optical density

Biomass is produced at the rate at which metabolites are drained from the model. Because $r_{\mathrm{BM}}$ was derived in Eq. (S11) in units of M/s, i.e. mol of precursor molecules per cell volume per time, we need to multiply it by the volume of a cell to get the change of the total biomass

$$\frac{\mathrm{d(total\ biomass)}}{\mathrm{d}t} = V_{\mathrm{cell}}\, r_{\mathrm{BM}}. \tag{S19}$$

In order to convert the biomass production rate $r_{\mathrm{BM}}$ to growth rate $\mu$, we need to take the total biomass concentration [BM] into account. This is easiest to see if we use the definition of growth rate, i.e. the relative change of biomass

$$\mu = \frac{1}{\mathrm{(total\ biomass)}} \frac{\mathrm{d(total\ biomass)}}{\mathrm{d}t} = \frac{r_{\mathrm{BM}}\, V_{\mathrm{cell}}}{\mathrm{(total\ biomass)}} = \frac{r_{\mathrm{BM}}}{[\mathrm{BM}]}. \tag{S20}$$

Because $r_{\mathrm{BM}}$ sets the time-scale of metabolic reactions, and $\mu$ sets the time-scale of protein turn-over, it is important to get these rates close to the biological reality to accurately describe both metabolite and protein dynamics. We discuss in detail how we derive these parameters in Section 3.

To compare biomass growth in experiments and theory, we derive optical density. Because optical density measures the total biomass, $\mathrm{OD600} \sim \mathrm{(total\ biomass)}$. Integrating Eq. (S20), we find

$$\mathrm{OD600}(t) = \mathrm{OD600}(0)\, e^{\int_0^t \mu(s)\, \mathrm{d}s}. \tag{S21}$$

### 3. Parameters of the model

#### 3.1. Motivation

Despite its simplicity, the model still contains a considerable number of parameters, including eleven reaction rates, five enzyme abundancies, four parameters for transcriptional regulation, four parameters describing Michaelis-Menten kinetics. In this section we estimate some of these parameters from the literature, and constrain the remaining parameters. We used the resulting parameter set in the majority of the paper, to describe all observed phenotypes of *E. coli*. We will describe how we obtained this parameter set first. Later in the paper, we screen the parameter space to get a sense of the possible solutions of the model, and trade-offs between phenotypes. For this purpose, we scanned a wide variety of parameters, which we will detail later in this section.

#### 3.2. Flux balance during steady state growth

The constraints for the parameters are two-fold: First, the system must be capable of growing in the designated conditions (i.e. glycolytic and gluconeogenic carbons), and second, the fluxes for each metabolite must be balanced to achieve steady state with efficient growth.

For steady state growth on a glycolytic carbon, we can derive from flux balance, that all fluxes going in, i.e. import of glycolytic carbon, must equal all outfluxes, i.e. drains of glycolytic and gluconeogenic carbons and biomass production,

$$r_{\emptyset\to\mathrm{GLY}} = r_{\mathrm{GLY}\to\emptyset} + r_{\mathrm{BM}} + 2r_{\mathrm{GNG}\to\emptyset}, \tag{S22}$$

with the factor 2 stemming from that fact that one glycolytic carbon is converted to two gluconeogenic carbons between FBP and PEP.

Next, we calculate the rates at the irreversible reaction of upper glycolysis/gluconeogenesis, $r_{\mathrm{up}}$. Here, the net flux through this reaction has equal the flux of biomass precursors branching off below upper glycolysis/gluconeogenesis and the lower drain,

$$r_{\mathrm{up}} = r_{\mathrm{GLY}\to\mathrm{FBP}} - r_{\mathrm{FBP}\to\mathrm{GLY}} = (\beta_{\mathrm{PEP}} + \beta_{\mathrm{GNG}})r_{\mathrm{BM}} + 2r_{\mathrm{GNG}\to\emptyset}. \tag{S23}$$

Because the net flux during steady state growth is given by flux balance, we have one parameter left to specify. We chose to $r_{\mathrm{GLY}\to\mathrm{FBP}} = (1+\psi_{\mathrm{up}})r_{\mathrm{up}}$ and $r_{\mathrm{FBP}\to\mathrm{GLY}} = \psi_{\mathrm{up}} r_{\mathrm{up}}$, such that $\psi_{\mathrm{up}}/(1+\psi_{\mathrm{up}})$ is the futile cycling ratio in the upper irreversible reaction. Using Eqs. (S3) & (S4), and the steady state metabolite concentrations [PEP*], [GLY*] and [FBP*], we derive

$$k_{\mathrm{cat}}^{\mathrm{pfk}}\,\phi_{\mathrm{pfk}}\frac{1}{1+K_{\mathrm{M}}^{\mathrm{pfk}}/[\mathrm{GLY}^*]} = (1+\psi_{\mathrm{up}})r_{\mathrm{up}} \tag{S24}$$

and

$$k_{\mathrm{cat}}^{\mathrm{fbp}}\,\phi_{\mathrm{fbp}}\frac{1}{1+K_{\mathrm{M}}^{\mathrm{fbp}}/[\mathrm{FBP}^*]} = \psi_{\mathrm{up}} r_{\mathrm{up}}. \tag{S25}$$

We will estimate Michaelis-Menten constants and substrate concentrations in steady state, which determine the fraction on the left-hand side, later in this section. $k_{\mathrm{cat}}^{\mathrm{pfk}}\,\phi_{\mathrm{pfk}}$ and $k_{\mathrm{cat}}^{\mathrm{fbp}}\,\phi_{\mathrm{fbp}}$ are effective parameters that will be calculated from the above equations. To calculate the model for the main figures we used $\psi_{\mathrm{up}} = 0.3\%$.

Analogously, we describe the irreversible reaction at lower glycolysis/gluconeogenesis, $r_{\mathrm{low}}$. The net flux through this reaction has equal the flux of biomass precursors branching off below lower glycolysis/gluconeogenesis and the lower drain,

$$r_{\mathrm{low}} = r_{\mathrm{PEP}\to\mathrm{GNG}} - r_{\mathrm{GNG}\to\mathrm{PEP}} = \beta_{\mathrm{GNG}} + 2r_{\mathrm{GNG}\to\emptyset}. \tag{S26}$$

As above, we choose $r_{\mathrm{PEP}\to\mathrm{GNG}} = (1+\psi_{\mathrm{low}})r_{\mathrm{low}}$ and $r_{\mathrm{GNG}\to\mathrm{PEP}} = \psi_{\mathrm{low}} r_{\mathrm{low}}$, such that $\psi_{\mathrm{low}}/(1+\psi_{\mathrm{low}})$ is the futile cycling ratio in the lower irreversible reaction. We then use Eqs. (S7) & (S8) to derive

$$k_{\mathrm{cat}}^{\mathrm{pyk}}\phi_{\mathrm{pyk}}\frac{1}{1+K_{\mathrm{M}}^{\mathrm{pyk}}/[\mathrm{PEP}^*]} = (1+\psi_{\mathrm{low}})r_{\mathrm{low}} \tag{S27}$$

and

$$k_{\mathrm{cat}}^{\mathrm{pck}}\phi_{\mathrm{pck}}\frac{1}{1+K_{\mathrm{M}}^{\mathrm{pck}}/[\mathrm{PEP}^*]} = \psi_{\mathrm{low}} r_{\mathrm{low}} \tag{S28}$$

which we use to calculate $k_{\mathrm{cat}}^{\mathrm{pyk}}\phi_{\mathrm{pyk}}$ and $k_{\mathrm{cat}}^{\mathrm{pck}}\phi_{\mathrm{pck}}$ as effective parameters. To calculate the model for the main figures we used $\psi_{\mathrm{low}} = 1.8\%$.

Lastly, we turn to the reversible reaction that connects FBP and PEP, Eqs. (S5) & (S6). This reaction is reversible and catalyzed by a single ‘effective’ enzyme, the super-enolase. This means, that both forward and backward reactions are proportional to the same protein abundance $\phi_{\text{eno}}$. In addition, the forward and backward reaction rates can theoretically not be chosen freely, but need to obey the thermodynamic constraint that the ratio is given by the change in Gibbs Free Energy $\Delta G$ between FBP and PEP $r_{\text{FBP}\to\text{2PEP}}/r_{\text{2PEP}\to\text{FBP}} = \exp(-\Delta G/RT)$. Here, $R$ is the gas constant and $T$ temperature. Plugging in Eqs. (S5) & (S6) we find

$$\exp(-\Delta G/RT) = \frac{r_{\text{FBP}\to\text{2PEP}}}{r_{\text{2PEP}\to\text{FBP}}} = \frac{k_{\text{cat}}^{\text{f}}\phi_{\text{eno}}[\text{FBP}]_{\text{gly}}}{k_{\text{cat}}^{\text{r}}\phi_{\text{eno}}[\text{PEP}]_{\text{gly}}{}^{2}} = \frac{k_{\text{cat}}^{\text{f}}[\text{FBP}]_{\text{gly}}}{k_{\text{cat}}^{\text{r}}[\text{PEP}]_{\text{gly}}{}^{2}} = \frac{\tilde{k}_{\text{cat}}^{\text{f}}}{\tilde{k}_{\text{cat}}^{\text{r}}} \quad \text{(S29)}$$

Where the subscript ‘gly’ refers to concentrations in glycolytic growth, and the tilde denotes that the rates were normalized to the glycolytic steady state. To understand the impact of the Gibbs Free Energy on fluxes in glycolysis, we calculate the ‘efficiency’ of the super-enolase and the relative net flux $E = (r_{\text{FBP}\to\text{2PEP}} - r_{\text{2PEP}\to\text{FBP}})/r_{\text{FBP}\to\text{2PEP}} = 1 - \exp(\Delta G/RT)$ . For $\Delta G = -7.79\text{kJ/mol}$ we obtain an efficiency $E = 95\%$.

In addition, the reversible reaction must obey flux balance, such that the net flux $r_{\text{super-eno}}$ suffices to feed biomass produced from precursors at PEP and TCA carbons, as well as the downstream drain,

$$r_{\text{super-eno}} = r_{\text{FBP}\to\text{PEP}} - r_{\text{PEP}\to\text{FBP}} = (\beta_{\text{PEP}} + \beta_{\text{GNG}})r_{\text{BM}} + 2r_{\text{GNG}\to\emptyset} = r_{\text{up}}. \quad \text{(S30)}$$

$$k_{\text{cat}}^{\text{f}}\phi_{\text{eno}}[\text{FBP}^{*}] = (E^{-1} - 1)r_{\text{super-eno}} \quad \text{(S31)}$$

and

$$k_{\text{cat}}^{\text{r}}\phi_{\text{eno}}[\text{PEP}^{*}]^{2} = E^{-1}r_{\text{super-eno}} \quad \text{(S32)}$$

*Table 1 Gibbs Free Energy (ΔG) change from FBP to PEP. Between FBP and PEP, metabolites are converted in a series of reactions by a series of distinct enzymes, shown in Box 1 (left). Changes of Gibbs Free Energy of reactions in between FBP and PEP, generated using eQuilibrator (Flamholz et al., 2012). When calculating the total change of Gibbs Free Energy, we need to take the stoichiometry into account, because one FBP yields two G3P. This results in a total change of Free Energy of* $\Delta G = \Delta G_{fba} + \Delta G_{tpi} + 2(\Delta G_{gap} + \Delta G_{pgk} + \Delta G_{gpm} + \Delta G_{eno})$.

| **Reaction** | **Enzyme** | **ΔG° in kJ/mol** |
|---|---|---|
| FBP <-> GP +G3P | fba | 22.05 |
| GP <-> G3P | tpi | 5.62 |
| G3P <-> BGP | gap | -1.83 |
| BGP <-> 3PG | pgk | -16.56 |
| 3PG <-> 2PG | gpm | 4.47 |
| 2PG <-> PEP | eno | -3.81 |
| FBP <-> PEP | 'super-eno' | -7.79 |

#### 3.3. Renormalization of model parameters

In addition to imposing constraints on the model, we introduce renormalizations of model parameters to reduce the number of effective parameters we need to use when calculating the model. In particular, we introduce relative Michaelis-Menten constants

$$\tilde{K}_{\mathrm{M}} = \frac{K_{\mathrm{M}}}{c^*}, \qquad \text{(S33)}$$

by dividing the Michaelis-Menten constant by the concentration of the respective metabolite in steady state $c^*$. By dividing catalytic rates by the glycolytic steady state concentrations, we introduce rescaled rates for the reversible reaction

$$\tilde{k}_{\mathrm{cat}}^{\mathrm{f}} = k_{\mathrm{cat}}^{\mathrm{f}} \phi_{\mathrm{eno}} [\mathrm{FBP}]_{\mathrm{gly}}, \qquad \text{(S34)}$$

$$\tilde{k}_{\mathrm{cat}}^{\mathrm{r}} = k_{\mathrm{cat}}^{\mathrm{r}} [\mathrm{PEP}]_{\mathrm{gly}}{}^2 \qquad \text{(S35)}$$

and analogously for all other rates.

#### 3.4. Absolute quantification of metabolite fluxes

In this subsection, we collect and derive biochemical parameters of the model. We start by estimating, the uptake rate of glycolytic carbon, $r_{\mathrm{BM}}$ , which determines the timescale of metabolic changes. For *E. coli* for a biomass density of 0.2 g dry weight per ml (Basan et al., 2015) growing on glucose at rate 0.9/h, the biomass production flux is $r_{\mathrm{BM}} = 0.2 \cdot 0.9\ \mathrm{g\ dwt/(h\ ml)}$. We can convert this rate to equivalents of mM glucose, using the biomass yield of glucose of 0.45 g glucose/g dry weight [3] and the molecular weight of 180 g/mol. In addition, we change units from hours to seconds to obtain

$$r_{\mathrm{BM}} = \frac{0.9 \cdot 0.2}{0.45 \cdot 3600 \cdot 180} \frac{\mathrm{h\ g\ (dwt)\ g\ (glucose)\ mol}}{\mathrm{s\ h\ ml\ g\ (dwt)\ g\ (glucose)}} = 0.62 \frac{\mathrm{mM}}{\mathrm{s}} \qquad \text{(S36)}$$

We can set the two drains such that they are small compared to the influx, e.g. $r_{\mathrm{GLY}\to\emptyset} = 0.1 r_{\emptyset\to\mathrm{GNG}}$ and $r_{\mathrm{GNG}\to\emptyset} = 0.05 r_{\emptyset\to\mathrm{GLY}}$. Using Eq. (S1), and the stochiometric conversion of one FBP to two gluconeogenic carbons, we calculate

$$\mathrm{V}_{\mathrm{max}}^{\mathrm{gly,up}} = \frac{r_{\mathrm{BM}}}{(1 - 0.1 - 0.025)} = 0.71 \frac{\mathrm{mM}}{\mathrm{s}} \qquad \text{(S37)}$$

Using Eqs. (S2) & (S9) we can calculate the drains relative to glycolytic steady state concentration of the respective carbons $\tilde{V}_{\mathrm{max}}^{\mathrm{gly,drain}} = V_{\mathrm{max}}^{\mathrm{gly,drain}} / [\mathrm{GLY}^*]$ and $\tilde{V}_{\mathrm{max}}^{\mathrm{gng,drain}} = V_{\mathrm{max}}^{\mathrm{gng,drain}} / [\mathrm{GNG}^*]$.

#### 3.5. Metabolite concentrations and Michaelis constants

Absolute concentrations of the metabolites are notoriously difficult to measure. In our work, we are guided by the metabolite concentrations measured in Ref. (Kochanowski et al., 2013a), see

Table 2, to set the steady state concentrations for growth on glucose. The absolute concentrations enter the model at the renormalized Michaelis-Menten constants, Eq. (S33).

*Table 2 Metabolite concentrations during steady state growth of Escherichia coli on glucose and acetate obtained from Ref. (Kochanowski et al., 2013a).*

| Metabolite | Growth on acetate | Growth on glucose |
|---|---|---|
| F6P (g/l) | 0.379 | 0.318 |
| FBP (g/l) | 0.149 | 1.380 |
| PEP (g/l) | 0.387 | 0.049 |
| F6P (mM) | 1.458 | 1.225 |
| FBP (mM) | 0.438 | 4.058 |
| PEP (mM) | 2.304 | 0.289 |

We obtained Michaelis-Menten parameters from the BRENDA database (www.brenda-enzymes.org), see Table 3.

*Table 3 Michaelis-Menten constants obtained from BRENDA database.*

| Enzyme | Michaelis-Menten constant (mM) |
|---|---|
| pfkA | 0.16 (Zheng and Kemp, 1995) |
| fbp | 0.015 (Donahue et al., 2000) |
| pyk | 0.083 (Berman and Cohn, 1970) |

We use metabolite concentrations and Michaelis constants to inform our choice of renormalized Michaelis constants. Since there is considerable uncertainty in metabolite concentrations, Michaelis constants and how to precisely transfer numbers to our coarse-grained model, we choose parameters in the vicinity of the experimental data that worked well in our model, see Table 4. In particular, for upper glycolysis, we choose $\tilde{K}_{\mathrm{M}} = 2$, which means that during glycolytic growth, the upper glycolytic enzyme is slightly undersaturated (rather than 0.16/0.318 = 0.5, which means slightly saturated). This choice ensures that sufficient flux can be catalyzed by upper glycolysis after nutrient shifts to glycolysis when glycolytic carbons build up. For upper gluconeogenesis we choose $\tilde{K}_{\mathrm{M}} = 0.05$, meaning that the enzyme is highly saturated in glycolytic growth (compared to 0.015/4.058 = 0.004). This choice ensures that futile cycling is low in glycolysis and that after shift to gluconeogenesis upper gluconeogenesis is close to saturation. For lower glycolysis we choose $\widetilde{K}_{\mathrm{M}} = 1$ (compared to 0.083/0.289 = 0.29). For lower gluconeogenesis we chose $\widetilde{K}_{\mathrm{M}} = 5$, such that the enzyme is undersaturated. This choice ensures that when nutrients are switched to gluconeogenesis that the enzyme becomes saturated. The chosen Michaelis constants ensure short lag times to glycolysis and a metabolic bottleneck in the shift to gluconeogenesis. Furthermore, they minimize wasteful proteome resources by keeping enzymes efficient.

*Table 4 Renormalized Michaelis parameters used in the model.*

| | Renormalized Michaelis constant $\widetilde{K}_{\mathrm{M}}$ |
|---|---|
| Upper glycolysis | 2 |
| Upper gluconeogenesis | 0.05 |
| Lower glycolysis | 1 |
| Lower gluconeogenesis | 5 |

#### 3.6. Allosteric parameters

While there is reliable information of which metabolites in central metabolism influence enzymatic turnover rates, translating this *in vitro* information to quantitative parameters in the model is difficult. In our model we thus chose to model allosteric interactions as power laws in Eqs. (S3), (S4), (S7) & (S8). The strength of these allosteric parameters determines the magnitude of futile cycling and recovery in both shifts. We randomly scanned the range of parameters to find a set of allosteric parameters that 1) produce low futile cycling in both glycolysis and gluconeogenesis and 2) yielded a lag of about 5h to gluconeogenesis and 1h to glycolysis, and found the set

Table 1, which we used throughout the paper.

*Table 5 Allosteric parameters used in the model.*

| Allosteric parameter | |
|---|---|
| $\alpha^{\mathrm{pfk}}$ | 0.5 |
| $\alpha^{\mathrm{fbp}}$ | 3 |
| $\alpha^{\mathrm{pyk}}$ | 0.95 |
| $\alpha^{\mathrm{pck}}$ | 0 |

##### 3.7. Biomass composition

Precursors for biomass are derived from metabolites at different stages in central metabolism, which is contained in our model in the 'biomass vector' $\beta$. From Ref. (Gerosa et al., 2015), we calculate that $\beta_{\mathrm{GLY}} = 20.9\%$ of the biomass flux is derived from upper glycolysis (F6P, G6P, E4P and R5P), $\beta_{\mathrm{PEP}} = 14.6\%$ from central part (G3P, 3PG and PEP) and $\beta_{\mathrm{GNG}} = 64.4\%$ from TCA carbons (acCoA, OAA, GLU, GLN, PYR and AKG).

### 5. Appendix Figures

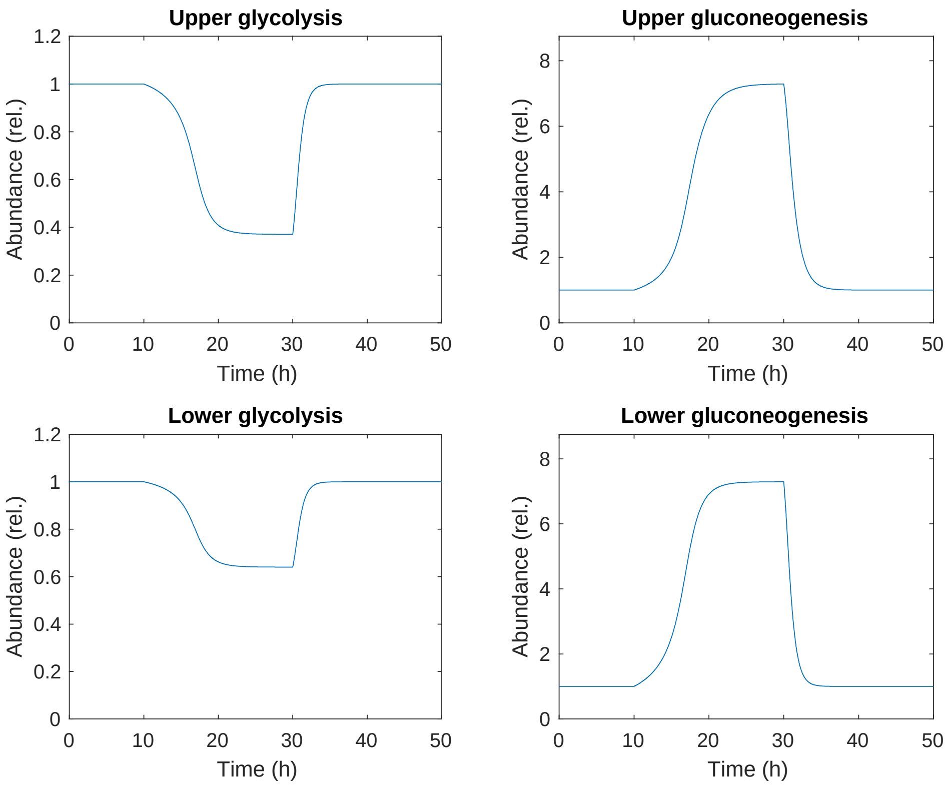

*Appendix Figure S1 Enzyme abundances relative to the glycolytic steady state at t=0h. Shift from glucose to acetate at t = 10h and acetate to glucose at t = 30h.*

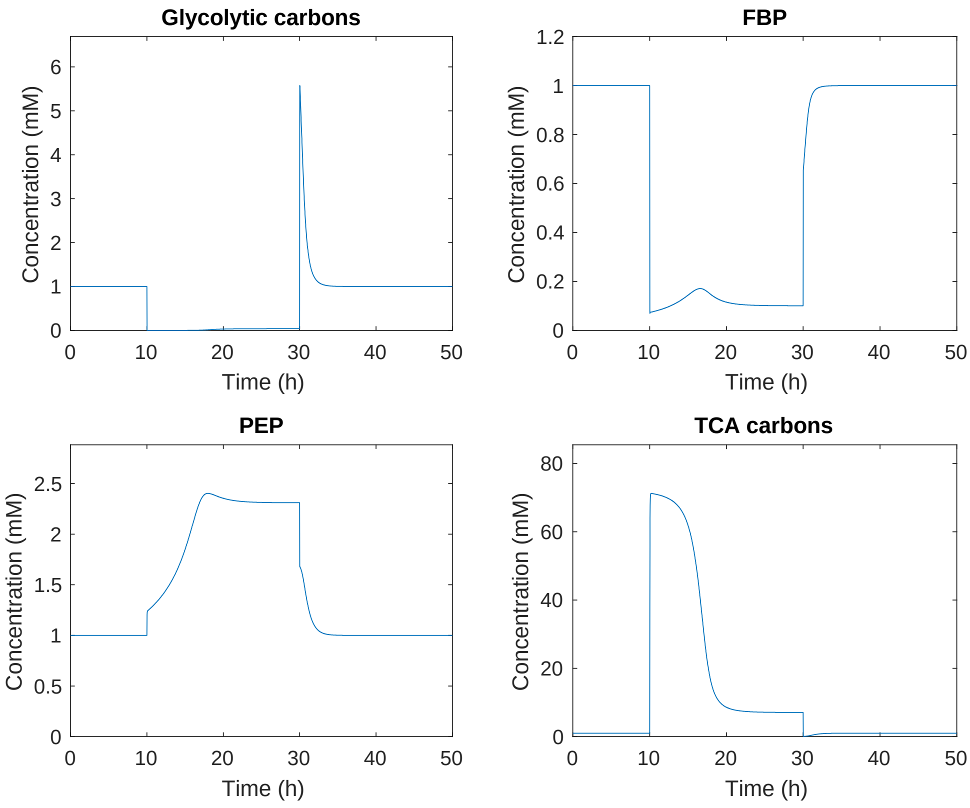

*Appendix Figure S2 Metabolite concentrations relative to the glycolytic steady state at t = 0h. Shift from glucose to acetate at t = 10h and acetate to glucose at t = 30h.*

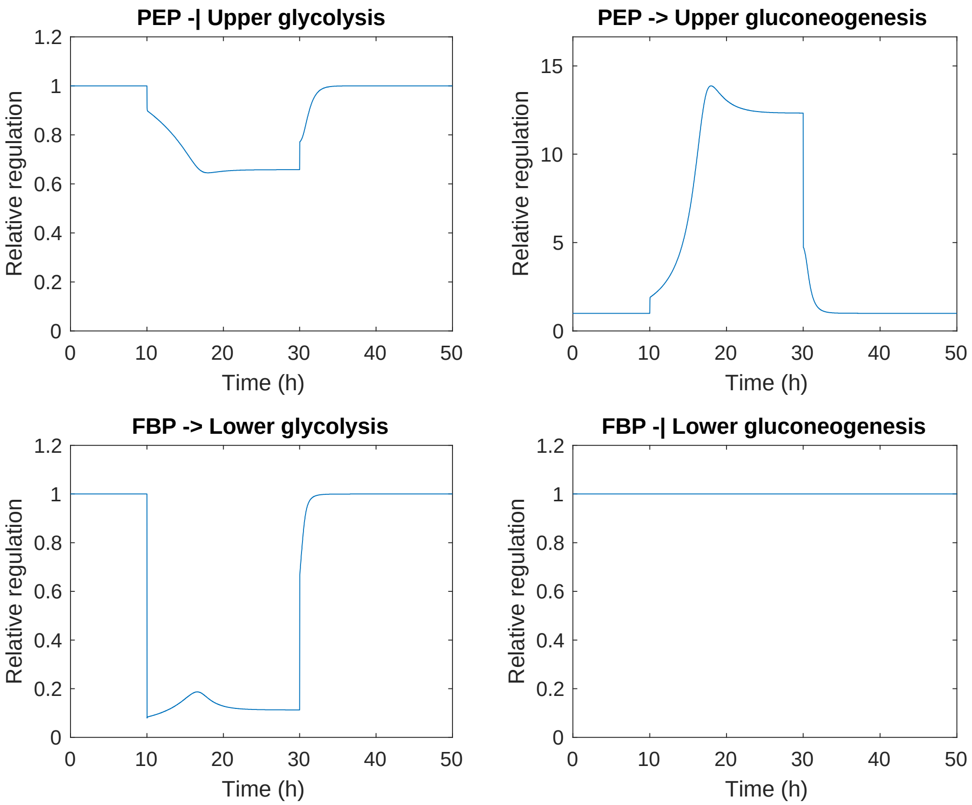

*Appendix Figure S3 Allosteric regulation relative to the glycolytic steady state t = 0h, defined as* $R_i = \left(c_i/c_i^{gly}\right)^{\pm\alpha_i}$*, where* $c_i$ *is the concentration of the regulatory metabolite (FBP or PEP) and* $\alpha_i$ *the strength of the allosteric regulation. The sign is positive if the enzyme is activated and negative for repression. Shift from glucose to acetate at t = 10h and acetate to glucose at t = 30h.*

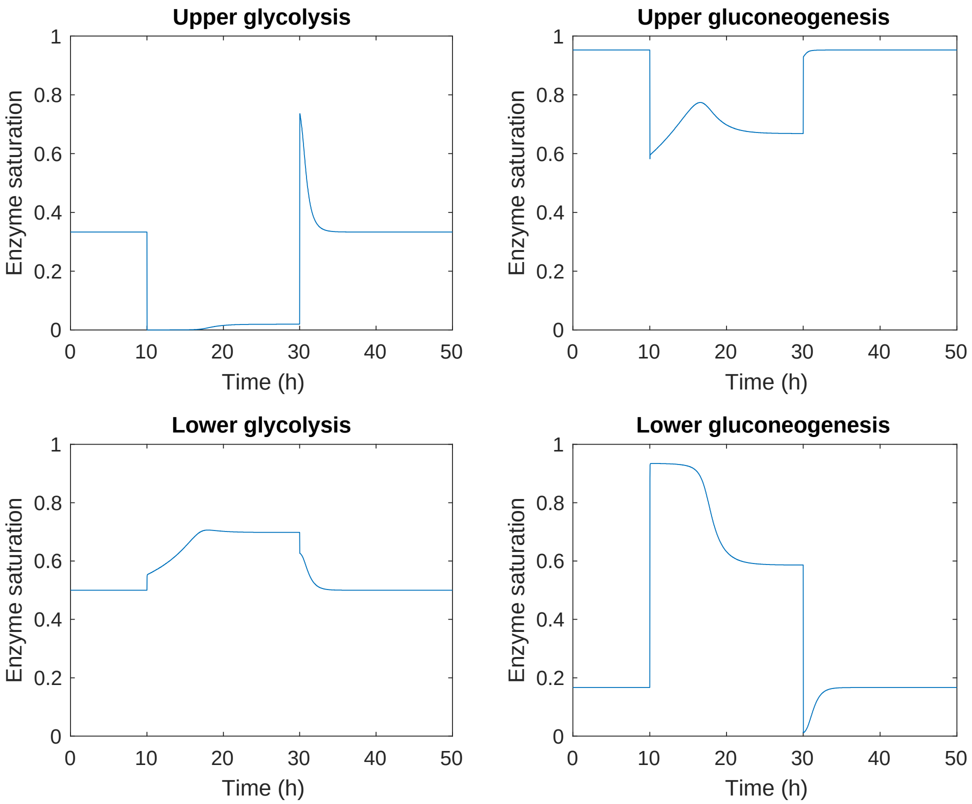

*Appendix Figure S4 Saturation of enzymes, defined as* $\frac{c_i}{c_i+K_M}$*, where* $c_i$ *is the substrate concentration and* $K_M$ *the Michaelis constant of the respective enzyme. Shift from glucose to acetate at t = 10h and acetate to glucose at t = 30h.*

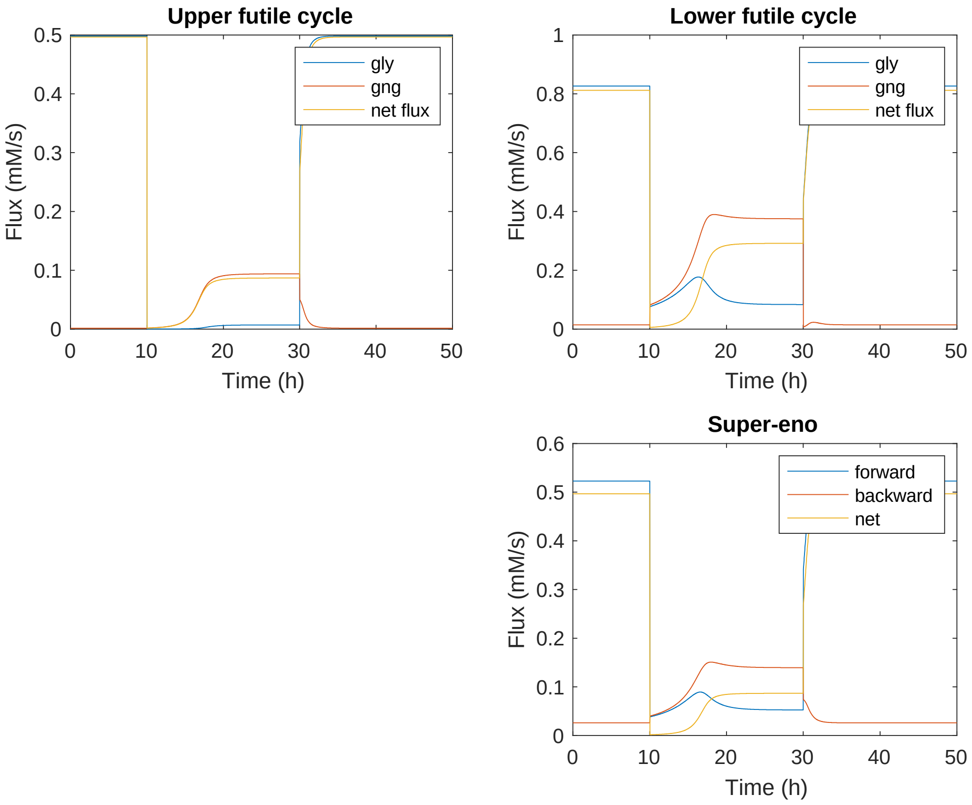

*Appendix Figure S5 Metabolic fluxes in the upper and lower reversible reactions, as well as the reversible 'super-eno' reactions. Shift from glucose to acetate at t = 10h and acetate to glucose at t = 30h.*

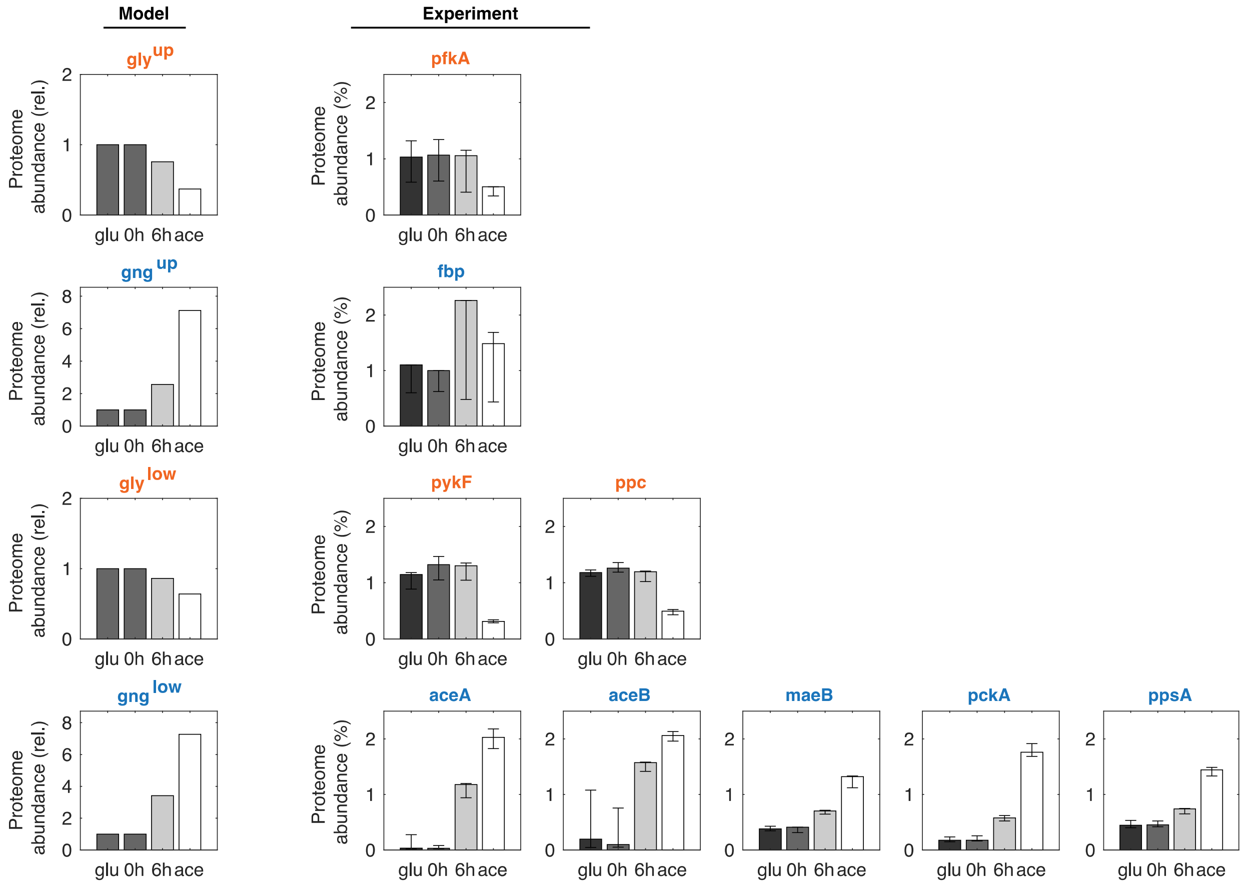

*Appendix Figure S6 Proteome fraction (model: relative to glucose steady state, experiment: fraction of total proteome) of glycolytic enzymes (orange) and gluconeogenic enzymes (blue) in a shift from glucose to acetate. Steady state on glucose: 'glu'. Steady state on acetate: 'ace'. Error bars show lower and upper quartiles. Proteome data from Ref. (Basan et al., 2020).*
